# Deep learning-based survival prediction using DNA methylation-derived 3D genomic information

**DOI:** 10.1101/2023.06.10.544450

**Authors:** Jeewon Yang, Dohoon Lee, Bonil Koo, Sun Kim

## Abstract

Three-dimensional (3D) genome states are closely related to cancer development. Nonetheless, the 3D genome information has not been clinically utilized to the best of our knowledge, due to the costly production of Hi-C data which is a manifest source of 3D genome information. Therefore, there is a need for a novel metric computable from a 3D genome-related data which is more easily accessible for the clinical utilization of 3D genome information. We here propose a method to extract 3D genome-aware epigenetic features from DNA methylation data and use these features for a deep learning-based survival prediction. These features are derived from the 3D genome structures which are rebuilt from the DNA methylation data in an individual level. The results showed that usage of 3D genome-aware features contributed to more accurate risk prediction across seven cancer types, suggesting the effectiveness of the knowledge about 3D genome structure embedded in these features. The deeper biological investigation revealed that altered DNA methylation level in risk-high group could be related to the anomalously activated genes involved in cancer-related pathways. Altogether, the risks predicted from 3D genome-aware epigenetic features showed its significance as a survival predictor in seven cancer types, along with its biological importance.

## Introduction

The organization of 3D genome plays an essential role in cell fate decision and the establishment of cell identity by providing spatial constraints to the action of transcription factors (TF)^1^. This implies that misconfigured 3D genome structure can activate aberrant transcriptional program, which may cause malignant transformation of cells and ultimately lead to cancer^2, 3^. The 3D genome structure is also known to drive oncogenic structural variations^4^, including driver fusion events by which proto-oncogenes hijack enhancers to facilitate their expression^5^. Altogether, it can be considered that the 3D genome organization underlies various aspects of cancer biology and its disruption can cause detrimental effects to the cell fate. However, to the best of our knowledge, there has been no study utilizing the quantified 3D genomic information in the cancer prognosis prediction. This results from two main reasons: (1) the low accessibility to the data capturing the 3D genome conformation of cancers, and (2) the lack of measures to evaluate the 3D genome perturbation quantitatively.

The most favored data representation for the snapshot of the 3D genome landscape has been contact matrices, or contact maps, produced by high-throughput chromosome conformation capture (Hi-C) experiments^6^, since they provide us with direct information about interplay between different genomic regions. In detail, contact maps show us how frequently two arbitrary genomic loci contact, which is a clue to estimate the strength of interaction between them. However, there exist critical limitations in producing contact matrices: the high cost and complex procedures of Hi-C experiments^7, 8^. Such challenges result in the limited availability to 3D genome contact maps. Therefore, an alternative data which is far more plentiful and also contains 3D genomic information is needed for the development of prognostic score.

Interestingly, it is discovered that 3D genome states can be also reconstructed from DNA methylation data^9^. Since the interaction between distant genomic loci in turn leads to the co-varying DNA methylation levels of CpGs within these loci, the correlative structure of DNA methylation profiles could reflect the 3D genome organization. Especially, distribution of A/B compartments, which are defined on multi-megabase scale and contrast with each other on the chromatin conformation^10–12^, were reconstructed from a set of DNA methylation data. Data produced by DNA methylation microarray techniques, including Illumina Infinium HumanMethylation450 (450K) BeadChip array, were used for this purpose^13^. The DNA methylation profiles are available for many cancer types in a patient-wise manner, contrary to Hi-C contact matrices, providing us with richer information.

Although the potential of DNA methylation data for 3D genome inference has been illuminated, conjecturing the individual 3D genome state has not been available yet. We developed a method for extracting DNA methylation-derived individual 3D genome information, henceforth called the 3D genome-aware epigenetic features, and predicted risk of failures. In detail, the differential structure of DNA methylation which approximates the correlative pattern was captured. Based on the observation that open sea CpG positions located distant from CpG islands showed the greatest predictive power in 3D genome reconstruction^9^, the same probes were used. We implemented a deep learning-based survival prediction model utilizing an existing method^14^. The risks were predicted by the model, using the 3D genome-aware epigenetic features as input. Afterwards, a pan-cancer survival analysis along with biological interpretation were conducted.

In this work, we propose a novel way of risk prediction using the 3D genome-aware epigenetic features. The pan-cancer analysis revealed the significantly different survival patterns between risk-high/low groups, suggesting the potential of predicted risks as an important prognostic predictor. The functional annotation on genes located in differentially methylated regions (DMRs) disclosed the involvement of DMR genes in many cancer-related pathways. One characteristic pathway was the retinoic acid (RA) signaling pathway, which is crucial for the developmental process^15^. The investigation on the chromatin states of DMRs showed that most DMRs have inactive or moderately activated states, implying the relationship between the altered DNA methylation level and anomalous activation of genes. To conclude, the 3D genome-reflective epigenetic features serve as important predictors of cancer prognosis and possess biological importance.

## Results

### BDM PC1s can approximate Hi-C PC1s

First, the validation on whether the BDM PC1s can reproduce Hi-C PC1s was conducted. The 450K DNA methylation data of different TCGA cohorts and stem cells from the Progenitor Cell Biology Consortium (PCBC) were used^16–18^, along with the PC1s from the raw Hi-C matrices^19, 20^. The compositions of TCGA and PCBC dataset is described in Table S1 and Table S2, respectively. The detailed procedures of processing raw Hi-C data are explained in the Supplementary information section **Hi-C data processing**.

Due to the large number of samples for each category (normal, tumor, and stem cells), the PC1s were averaged from 10 random samples of the same category. For a fair comparison, BDM PC1 and Hi-C PC1 from the same category and tissue type were paired (Table S3-S5), and evaluated by PCC. The results showed that BDM PC1s can reproduce Hi-C PC1s to a dependable extent (PCC over 0.5) in most cases, with the highest performance observed in the reproduction of cancer cell’s Hi-C PC1s (Fig. 1 and Fig. S3).

**Figure 1.**
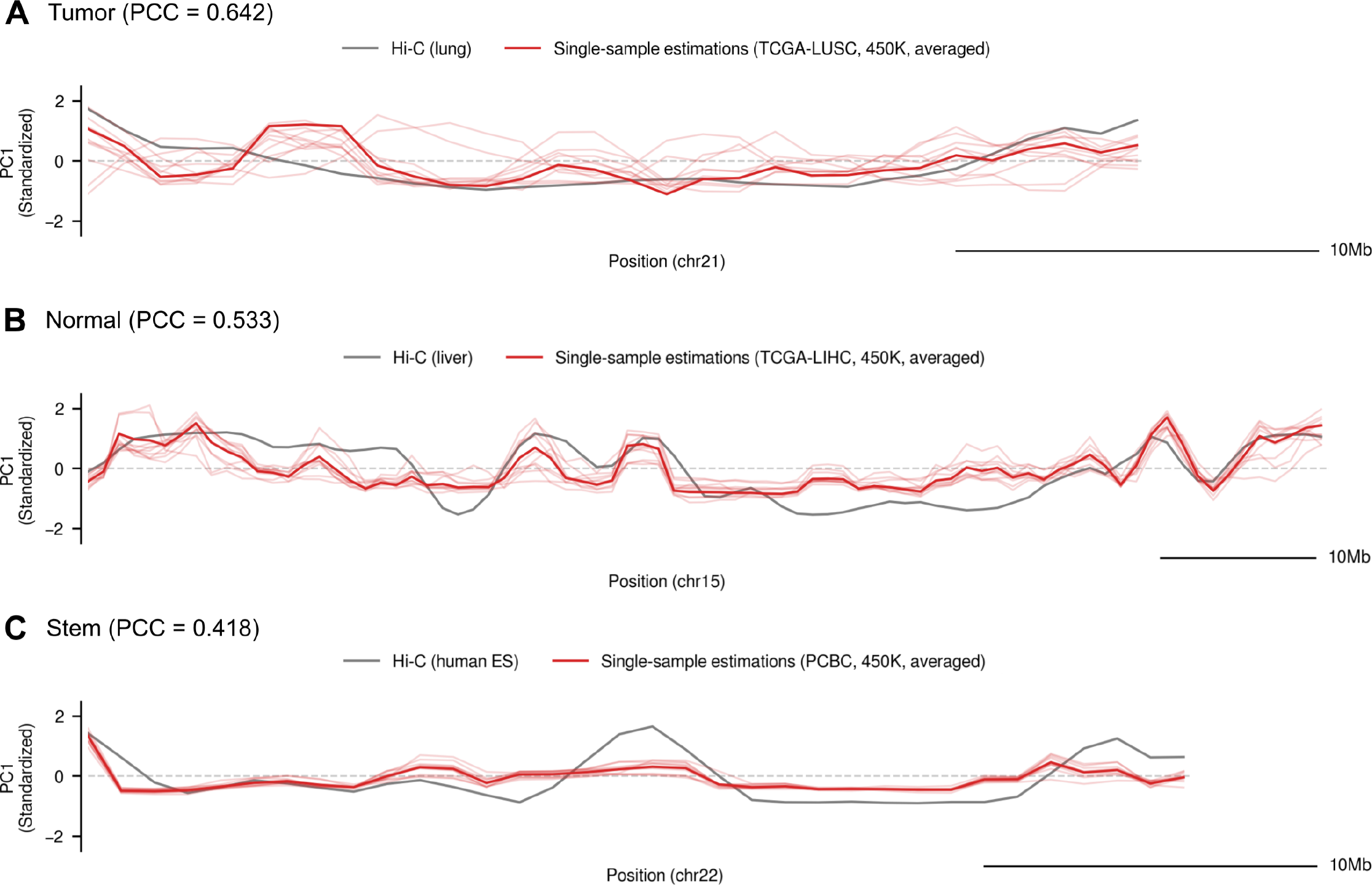
Reproduction of Hi-C PC1s from BDM PC1s. The dark red graph depicts the averaged PC1s from 10 samples. The light red graph illustrates individual BDM PC1. The gray graph shows the Hi-C PC1. Brief explanations are written at the top center of each subplot. The category of PC1 (tumor, normal, or stem) and the Pearson correlation coefficient (PCC) between the averaged BDM PC1 and Hi-C PC1, are written at the top left of each subplot. (A) BDM PC1 from tumor samples (TCGA-LUSC, chr21) and Hi-C PC1 from lung squamous cell carcinoma. (B) BDM PC1 from normal samples (TCGA-LIHC, chr15) and Hi-C PC1 from normal lung cells. (C) BDM PC1 from stem cells (PCBC, chr22) and Hi-C PC1s from human embryonic stem cells.

Considering that our approach is single-sample based, the comparison between individual PC1s were also conducted. As a result, the highest PCC among all categories increased to 0.750 (Fig. S4A) compared to the former case. This implies that the 3D genome information embedded in Hi-C PC1 can be better reproduced by using individual BDM PC1s rather than the averaged ones. It is conceived that using averaged BDM PC1 might lower the performance, as the well-reproduced individual BDM PC1 can be diluted by averaging it with other samples’ PC1s.

### BDMs and BDM PC1s capture innate differences between tumor and normal groups

To investigate whether BDM contains cancer-related 3D genomic information, which would necessarily differ between tumor and normal groups, the BDM heatmaps from two groups were compared. As a result, a recognizable patchiness was detected only in tumor groups across many cohorts (Fig. S5). Therefore, the BDMs are conceived to contain information regarding the inherent dissimilarity between tumor and normal groups. This tendency was also observed in BDM PC1s. The shape of PC1 plots were distinctly different between the two groups, across different cohorts (Fig. S6).

### BDM PC1s are tissue type-specific

In addition to capturing the tumor-normal difference, we conceived that BDM PC1s should be tissue type-specific for a pan-cancer clinical use. For this purpose, the BDM PC1s were grouped into different pairs, the homogeneous pair consisting of two PC1s from the same cohort and the heterogeneous pair whose PC1s originate from different cohorts, and PCC values were examined for each pair. As a result, PCC values from the homogeneous pairs were larger than those from the heterogeneous pairs (Fig. S7). Furthermore, among the heterogeneous pairs, those consisting of PC1s from the cohorts related to the same tissue type (e.g., KIRP and KIRC) showed bigger PCC values than the other pairs. These results imply that BDM PC1s encompass the tissue type-specific information.

### Utilizing 3D genome-aware epigenetic features helps predicting risk of failure

After the characteristics of BDM were probed, a one-dimensional vector consisting of 3D genome-aware epigenetic features and survival-related features was constructed to be used as an input feature for the deep learning model. To validate the usefulness of 3D genome-aware epigenetic features, two baseline scenarios were also investigated: (1) using no epigenetic feature (using only age and gender), and (2) using age, gender, and the 3D genome-unaware epigenetic feature (the average of open sea DNA methylation level) as input. For the evaluation metric, the C-index was used.

After the risk prediction, patients from each cohort were divided into risk-high/low groups, thresholded by the median risk value of the cohort. Subsequently, log-rank test was conducted to verify whether the predicted risk significantly influences the survival pattern. We conceived the results to be significant if both the validation and test c-index were greater than 0.65 and the log-rank test p-value was under 0.05. Consequently, significant results were observed in seven cohorts, suggesting the usability of the predicted risk as an important prognostic predictor (Table 1). The results from the log-rank tests are described in Fig. 2 and Fig. S8.

**Figure 2.**
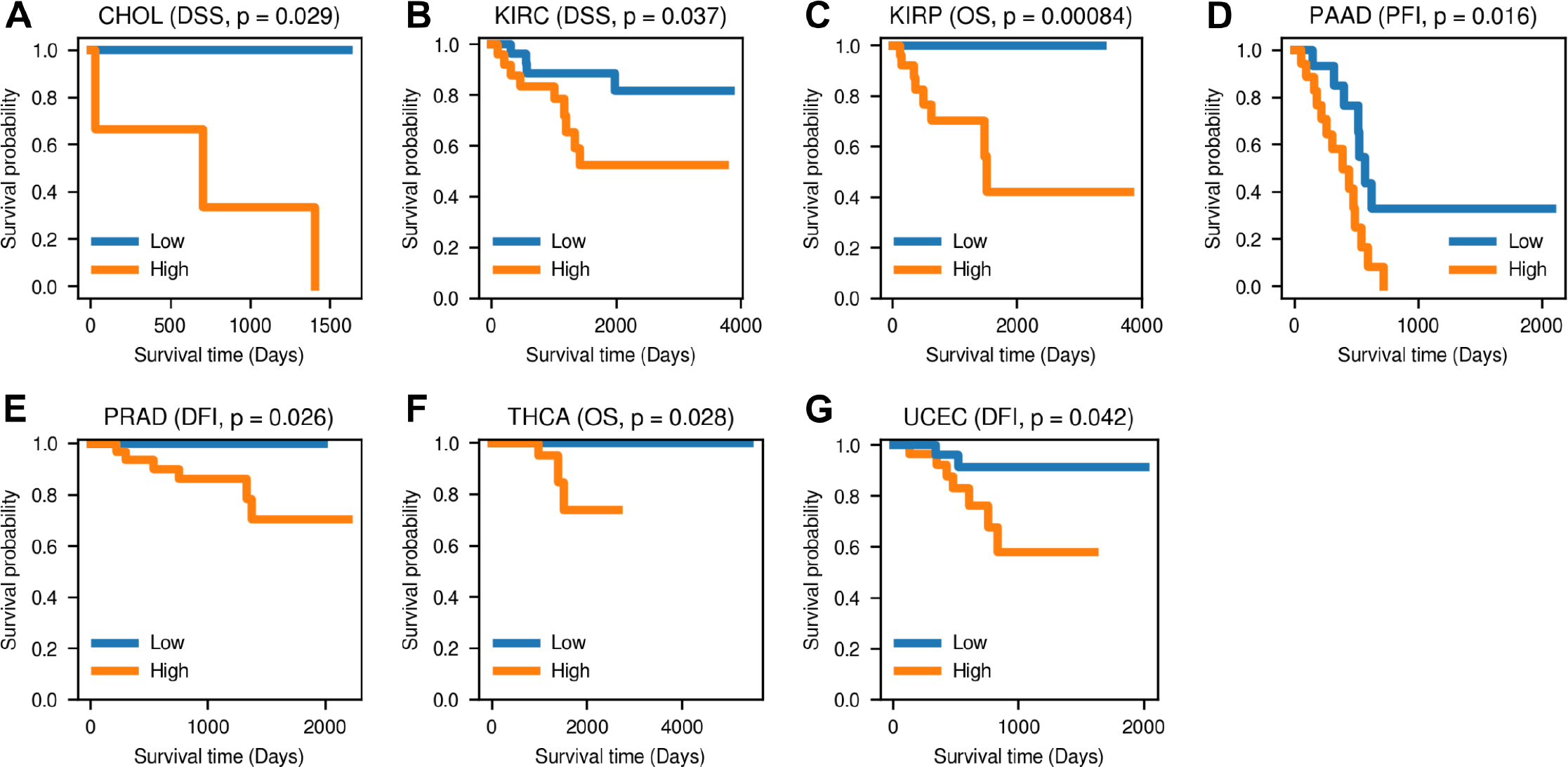
Log-rank test results based on the risks predicted by the feedforward neural network. One result per cohort is plotted. The other significant results are presented in Fig. S8. OS, DSS, DFI, and PFI stands for the overall survival, disease-specific survival, disease-free interval, and the progression-free interval, respectively. The name of the survival event and the log-rank test p-value are written in the parenthesis in the top center. (A) CHOL (DSS), (B) KIRC (DSS), (C) KIRP (OS), (D) PAAD (PFI), (E) PRAD (DFI), (F) THCA (OS) and (G) UCEC (DFI).

**Table 1.** Risk prediction results from using the 3D genome-aware epigenetic features along with age and gender. The best performances are highlighted in bold.

| Cohort | Test set c-index | Log-rank p-value | Event |
| --- | --- | --- | --- |
| CHOL | 0.750 | 0.029 | DSS |
| KIRC | 0.742 | 0.049 | DSS |
| KIRC | 0.705 | 0.045 | OS |
| KIRP | 0.841 | **0.001** | OS |
| KIRP | **0.959** | 0.024 | DSS |
| KIRP | 0.902 | 0.004 | DFI |
| KIRP | 0.839 | 0.040 | PFI |
| PAAD | 0.688 | 0.016 | PFI |
| PRAD | 0.739 | 0.026 | DFI |
| THCA | 0.742 | 0.028 | OS |
| UCEC | 0.654 | 0.038 | DSS |
| UCEC | 0.669 | 0.042 | DFI |

Following the log-rank test, Cox regression was also carried out. The results suggested that predicted risks are more important than other covariates in survival prediction (Fig. S9). Consistent with the results from the log-rank test, high risk had negative impact on the survival.

The results from two baseline cases showed that the survival prediction performance declines by excluding the 3D genomeaware epigenetic features from the input (Table S6; **BS1** and **BS2**). Additionally, both baseline scenarios did not show good performance for events other than OS. Considering that age is a very powerful predictor of OS, absence of 3D genome-reflective features might have caused the model to focus more on age, thereby making the model perform well only on OS.

### The model shows robust performance on external datasets

To validate the robust performance of the model on a dataset from other platform, the model was evaluated using GSE103659 (GEO accession GSE103659^21^). Because GSE103659 consisted of the glioblastoma (GBM) patients, the model performances on this dataset and TCGA-GBM were compared. The results showed that predicted risks were significant survival predictors for both datasets (Table S7 and Fig. S10). Taking into account that there was lack of information in GSE103659 compared to TCGA-GBM (the absence of gender data, and the smaller number of survival events with available data), it is suggested that although more significant results were observed in TCGA-GBM compared to GSE103659 (Table S7), the performance gap of that extent is at an acceptable level to consider. Consistently, the Cox regression results also demonstrated that predicted risk is a significant survival predictor in both TCGA-GBM and GSE103659 (Fig. S11).

### Functional annotation of genes in DMR

The functional annotation on genes located in DMR, defined by the predicted risks, revealed that many of them are involved in the RA signaling pathway. RA is known to bind to its receptor and induce conformation changes into the euchromatin structure, thereby provoking the transcription of target genes and playing an important role during the developmental process^15, 22^. Furthermore, RA is known to modulate the cell fate decisions, including differentiation, apoptosis, or stemness of a cell, in a cell-type specific manner^23^. Therefore, it can be presumed that the open sea CpG positions can epigenetically control the important regulators of development and cell fate and influence the stemness of cells, and such information is embedded in the 3D genome-aware features. Other cancer-related pathways, including gas transport, cell-cell adhesion, mitotic cell cycle, and gamma-aminobutyric acid (GABA) signaling pathways were also enriched. The thorough explanation on these results are provided in Supplementary information section **Functional annotation of DMR genes**. All results are described in Fig. 3 and Table S8.

**Figure 3.**
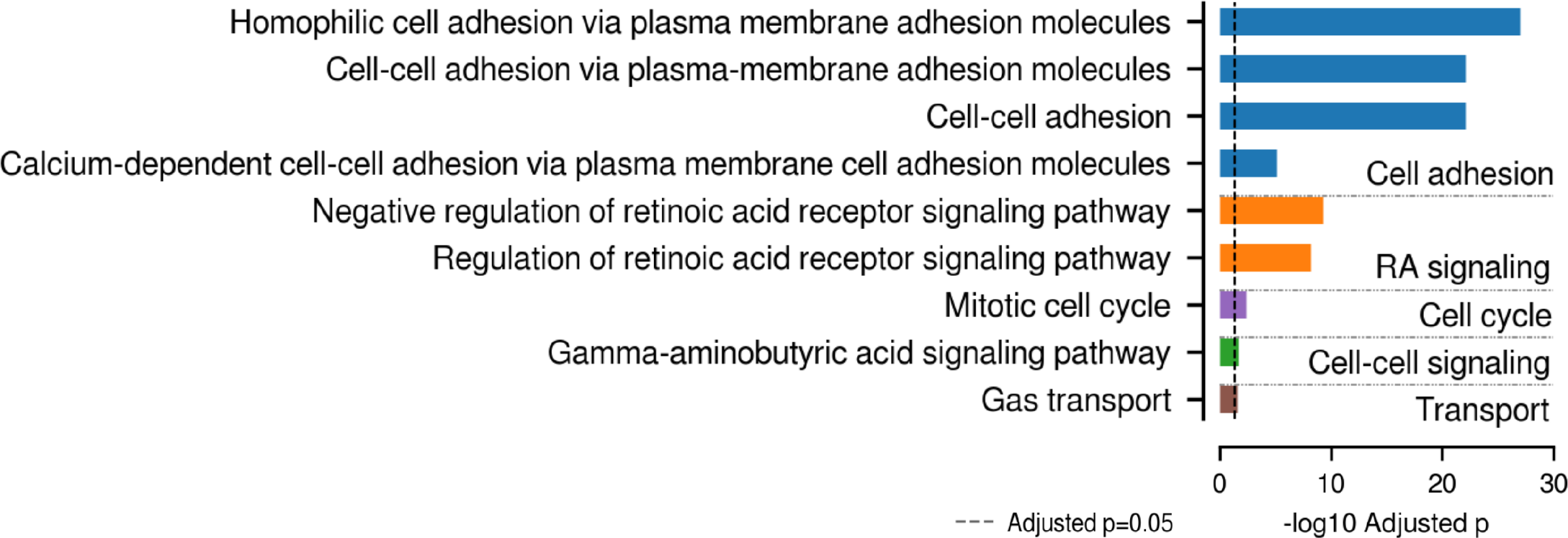
*−* log_10_(adjusted p-value) of each Gene Ontology (GO) term enriched by the functional annotation using genes in the differentially methylated regions (DMR) as input. GO terms from each category are split by the horizontal gray line, with the name of the category written at the right of the bars. The black vertical line demarcates *−* log_10_ 0.05.

### Inactive chromatin states take major proportion in DMRs

Using the chromatin state data^24, 25^, the proportion of each chromatin state in DMR was analyzed to figure out which states are the most influenced by cancer-related 3D genome aberrations. Among all cohorts in Table 1, the one with available chromatin state data was used (TCGA-PAAD). As a result, **Quiescent/Low** accounted for the largest proportion, followed by **Weak transcription** and **Heterochromatin** (Fig. S12). Among these dominant chromatin states, **Quiescent/Low** and **Heterochromatin** belong to the inactive states in normal cells. Additionally, **Weak transcription** is a mildly activated state. It is thus conceived that DMR genes could be abnormally influenced by the hypomethylation on open sea CpG positions accompanied with the progression of cancer.

## Discussion

Although the close relation between 3D genome and cancer development has been observed^4^, a prognostic metric which takes this into consideration has not been developed. The main reason for this is the costly production of Hi-C data, a manifest source of the 3D genome information, resulting in the low availability of Hi-C data^7^. Inspired by the rediscovery concerning the potential of 450K DNA methylation data to rebuild the 3D genome information^9^, the 3D genome-aware epigenetic features were derived from the DNA methylation data. Using these features, the risk of failure was predicted by a feedforward neural network. The predicted risk turned out to be a significant predictor of survival across many cancer types. One noteworthy result is that excluding the 3D genome-aware features from the input decreased the model performance. It can be posited that usage of 3D genome-aware features enables a knowledge-guided risk prediction, resulting in the more accurate prediction of cancer prognosis. Furthermore, the functional analyses showed that genes in DMR, defined by the risk values, participate in many cancer-related pathways including cell adhesion, RA signaling pathway, and mitotic cell cycle. And a comprehensive examination on the chromatin states of DMRs revealed that inactive or mildly activated states are dominant in these regions. From these results, it is postulated that the altered DNA methylation level in risk-high group is related to perturbed cancer-related pathways and abnormally activated genes. After thorough consideration, we conclude that 3D genome landscape inferred from DNA methylation data, which could be indicative of the aberrantly activated cancer-related genes and pathways, guides more accurate prediction of the cancer prognosis.

## Methods

### Task design and approach

The overall approach consists of two major parts: feature engineering and a deep learning-based survival prediction. After the features encompassing the 3D genome information are extracted from DNA methylation data, they are used as input for the pan-cancer survival prediction. The key concepts are described in the paragraphs below.

1. Inference of individual 3D genome enables quantifying 3D genome state of each cancer patient, increasing the clinical usability.
2. The inferred 3D genome structures are represented as vectors (the first principal components; PC1s), enabling the measurement of dissimilarities between two distinct individual 3D genome states.
3. The representations of normal or stem cells’ 3D genome structures, or **stem/normal references**, are constructed by averaging the 3D genome states of multiple samples.
4. The dissimilarity of each individual 3D genome state from normalness or stemness is quantified by measuring the distances between the individual 3D genome state and the reference (**stem/normal distance**). Using these distances, the **stem closeness** of each sample is measured.
5. We investigated three scenarios of risk prediction: using the 3D genome-aware, 3D genome-unaware, or no epigenetic features for a deep learning-based risk prediction.

Our study suggests that the 3D genome organization can be inferred as PC1 vectors from the DNA methylation levels of open sea CpG positions, and they contribute to more accurate risk prediction of cancer patients. Actually, representing the differential structure of DNA methylation states as PC1s has been widely used to illustrate the chromatin conformation^26, 27^. These PC1s stand for the inferred 3D genome states, whose entries identify the A/B compartment of the corresponding genomic bins. Using these vectors, dissimilarities between distinct individual 3D genome structures can be quantified and 3D genome-aware features can be extracted.

Fig. 4 describes the overview of the whole pipeline. Different from an existing method^9^, which infers a single tissue-type specific 3D genome state from multiple samples (Fig. S1A), the 3D genome structure is rebuilt from each individual (Fig. 4A and Fig. S1B). The dissimilarities between individual 3D genome states are measured by the distance between PC1s. Utilizing the normal/stem references, stem closeness of each individual is examined (Fig. 4B-C). These 3D genome-aware epigenetic features were concatenated along with age and gender, and used as the input for the risk prediction by a feedforward neural network (Fig. 4D). Following the pan-cancer survival analysis, functional analysis on DMRs were conducted to investigate the biological importance of the predicted risks. The more quantitative explanation on extracting 3D genome-aware epigenetic features are provided in Fig. S2.

**Figure 4.**
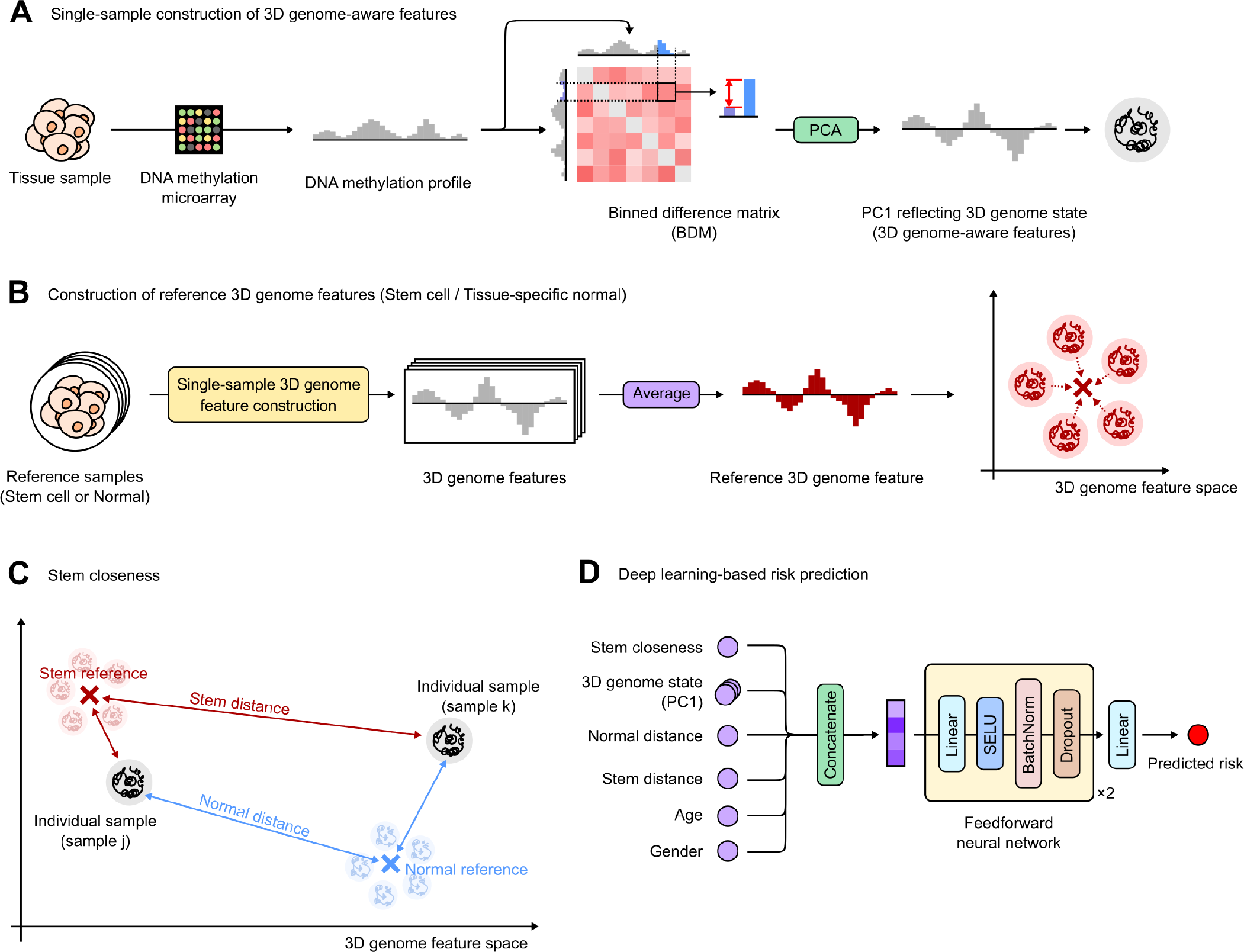
Overview of the whole pipeline. (A) Single-sample inference of 3D genome structure. The binned difference matrix (BDM) is constructed from the 450K DNA methylation data, and the PC1 of BDM is extracted. The BDM PC1s are reflective of the 3D genome structures. (B) Reference 3D genome states are constructed by averaging the BDM PC1s from multiple normal cells of the same tissue type or the stem cells. The resulting vectors are used as normal/stem references, representing the tissue type-specific normalness or stemness, respectively. (C) Each reconstructed 3D genome state is represented into a space. Distances from individual states to stem/normal references are measured. The blue and red arrows represent normal and stem distances, respectively. Between two depicted samples *j* and *k*, sample *j* has shorter stem distance and longer normal distance. Therefore, sample *j* has higher stem closeness. The more detailed procedures for quantifying stem closeness is explained in the supplementary information sections **Extracting 3D genome-aware epigenetic features from BDM** and **Figuring out the optimal stem closeness for each cancer type**. (D) The 3D genome-aware epigenetic features and survival-related features (age and gender) are concatenated for each sample. The feedforward neural network predicts risk of failures using this input feature.

### Deriving 3D genome-aware epigenetic features in an individual-level

Two classes of large genomic compartments with contrasting characteristics, namely euchromatic A and heterochromatic B compartments, constitute the top-level hierarchy of the 3D genome organization^28^. It is widely known that this compartmentalization can be inferred through the analysis of high-throughput measurements of genomic contacts, owing to the stark scarcity of inter-compartment contacts compared to intra-compartment contacts. Specifically, this correlative structure of 3D genome is manifested as a broad plaid pattern throughput Hi-C contact frequency matrix, therefore the PC1 values of normalized Hi-C matrix. The Hi-C PC1s, capturing the largest variability of the Hi-C matrix, reflect the compartmentalization states^20, 29, 30^.

Inspired by the previous observation of co-varying DNA methylation levels across genomic regions^9^, we thought that a matrix representing the absolute difference in DNA methylation levels between two arbitrary genomic bins may also reflect the correlative structure of the 3D genome. In other words, the rationale behind this matrix, hereafter called binned difference matrix (**BDM**), is that we expect smaller differences in methylation levels between the two genomic regions under spatial association (Supplementary information section **Constructing binned difference matrix (BDM)** and Fig. S2A).

### Investigating the characteristics of BDM

To utilize the BDMs for devising a prognostic score, we presumed that there are several criteria BDM should satisfy: (1) containing enough 3D genome information, (2) showing distinct differences between tumor and normal group, and (3) tissue-type specificity. Several experiments were conducted to verify whether BDMs comply with these requirements. To examine whether BDMs encompass the 3D genome information, the Pearson Correlation Coefficient (PCC) between BDM PC1s and Hi-C PC1s were measured. In addition, BDMs from tumor and normal groups were visualized into heatmaps to scrutinize the inter-group difference. Lastly, all-pairwise PCC between the averaged BDM PC1s were evaluated to ascertain that PCC values from homogeneous pairs consisting of the BDM PC1s from the same TCGA cohort is bigger than those from all the other cases.

### Devising a prognostic score from the BDM PC1s

The main assumption is that BDM PC1 reflects individual 3D genome structure, which allows for measuring the differences between distinct 3D genome states as the distances between vectors. Based on this concept, the stem- and normal-likeness of each cell were assessed using **stem/normal reference** (Fig. 4B and Fig. S2B). The resulting distances between each BDM PC1 and stem/normal reference, since named **stem/normal distance**, represent dissimilarity of each sample to stem cells’ or normal cells’ states (Fig. 4C and Fig. S2C). We explored two possible distance metrics: the euclidean distance and the inverse of cosine similarity. In case of the latter, a pseudocount (10*^−^*^1^^5^) was added to the similarity before the inversion to prevent division by zero. The biological meaning of each distance implies that a stem-like cell would have high normal distance and small stem distance. Our prognostic score, quantifying stem closeness of a cell, was designed based on this idea (Eq. 1). In Eq. 1, *d_n_* and *d_s_* denote normal and stem distance, respectively.

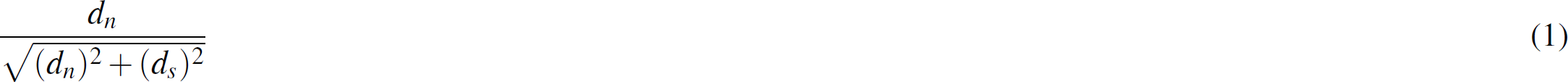

The more detailed procedures for extracting the 3D genome-aware features from BDM, along with the underlying rationale, are explained in the supplementary information sections **Extracting 3D genome-aware epigenetic features from BDM** and **Figuring out the optimal stem closeness for each cancer type**.

### Risk prediction using a feedforward neural network and 3D genome-aware epigenetic features

Provided that the aforementioned 3D genome-aware epigenetic features encompass cancer-related 3D genomic information, it was postulated that utilizing these features would bring about the outperforming results of survival prediction, compared to the baseline cases not using these features. As for baselines, we probed two scenarios: (1) using age and gender for survival predictors without any epigenetic features, and (2) using age and gender along with the 3D genome-unaware epigenetic feature (the average DNA methylation level of open sea CpG positions).

A feedforward neural network introduced by an existing paper^14^ was used to predict the risk of failure (Fig. 4D), consisting of two hidden layers with 128 hidden nodes in each layer. The average negative log partial likelihood was used as a loss function during training (Eq. 2). In Eq. 2, *N_E_*_=1_ denotes the number of patients from whom the event was observed, inline is a log-risk function estimated by the neural network, *i* and *j* are indices of patients, *R*(*T_i_*) is a set of patients who are at risk of failure at time *T_i_*, and *λ* is a L2 regularization coefficient.

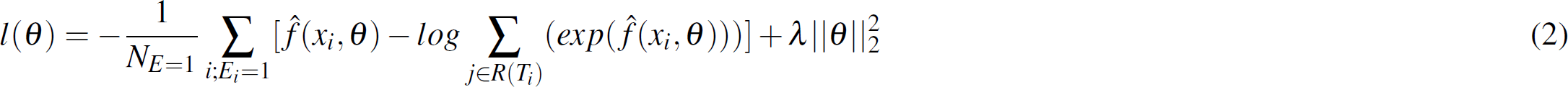

For the activation function and gradient descent algorithm, the scaled exponential linear units (SELU) and stochastic gradient descent (SGD) with nesterov momentum (momentum factor: 0.9) were used, respectively. To prevent overfitting, early stopping with patience 10, L2 regularization with coefficient 10, dropout with probability 0.4, and batch normalization were utilized. In addition, the time-based learning rate decay was employed to reduce the learning rate every epoch.

The performance of the model was evaluated by the Concordance Index (C-index) using lifelines^31^, for the four respective survival events: Overall Survival (OS), Disease-Specific Survival (DSS), Disease-Free Interval (DFI), and Progression-Free Interval (PFI). Given that C-index evaluates the degree of accordance between the predicted and ground-truth survival, the greater C-index was considered to be reflect the higher model performance. The dataset was constructed separately for each survival event in each cohort. After the samples without unavailable survival data were excluded, the remaining samples were randomly split into training, validation, and test datasets according to the ratio of 6:2:2.

### Survival analyses based on predicted risk

Following the risk prediction, log-rank test and Cox regression were conducted to verify the significance of the estimated risk as a prognostic predictor. For the log-rank test, patients from each cancer type were stratified into risk-high/low groups, thresholded by the median risk value. In case of the Cox regression, four covariates were used: age, gender, the average open sea DNA methylation level, and predicted risk.

### Functional analyses

Taking into account that risks are predicted from 3D genome-aware epigenetic features, we surmised that difference between the risk-high/low groups emanates from the different level of DNA methylation on the open sea CpG positions. Furthermore, provided that these DNA methylation levels hold cancer-related 3D genome information, the scrutinization on the DMR between the risk-high/low groups could provide the biological interpretation on how these groups differ. Based on such rationale, functional annotation on the DMRs defined by the predicted risks were conducted, following the processes explained in the paragraphs below.

First, we conducted functional annotation on DMR genes. DMRs were defined as genomic bins which are substantially hypomethylated in risk-high group compared to the risk-low group. The functional annotation on all genes located in DMRs was performed using a python package GSEApy^32^, based on the gene set ‘GO Biological Process 2015’^33, 34^.

Second, the chromatin states in DMR were analyzed. Chromatin states, the epigenetic annotation describing the functions of noncoding genomic regions, are known to possess the 3D genome information^28, 35^. The proportion of each chromatin state in the DMR was assessed to examine which states are the most influenced by the altered 3D genome structure.

## Code availability

All source codes are available at the following GitHub repository (https://github.com/jwyang21/3D-genome-risk-prediction).

## Acknowledgements

This work was supported by the Bio & Medical Technology Development Program of the National Research Foundation (NRF) and the Institute of Information & communications Technology Planning & Evaluation (IITP) grant funded by the Korea government (MSIT) [NO.2021-0-01343, Artificial Intelligence Graduate School Program (Seoul National University)].

## Author contributions

J.Y. and D.L. conceptualized the overall methodology and designed experiments. J.Y., D.L. and B.K. curated data. J.Y. conducted the experiments. J.Y. and D.L. wrote original draft for manuscript. J.Y., B.K., D.L., and S.K. analyzed the experiment results and reviewed the manuscript.

## Supplementary information

### Constructing binned difference matrix (BDM)

This section describes the detailed pipeline for constructing BDM. The main concept of BDM is to capture the differential structure of DNA methylation level among different genomic regions. Therefore, a representative value which stands for the overall DNA methylation level for each region should be defined so that the difference between these values can be measured. For this purpose, each autosome is split into 1Mb-sized bins. Then, the median value among all DNA methylation levels of open sea CpG probes belonging to each genomic bin is computed Fig. S2A). This was based on another main concept of BDM, which is to use distant or open sea CpG probes only, since they were known to have superior predictive power over CpG probes of other regions (island, shelf and shore) in terms of inferring 3D genome from DNA methylation level^9^. Then, the absolute difference between these median values are calculated for all possible pairs of bins. The resulting values are used as entries of BDM of this chromosome. The row and column indices of BDM are consistent with the bin indices from which the difference value resulted. This process is repeated for all autosomes. The explained processes are depicted in Fig. S2A.) For convenience, BDM construction of an arbitrary chromosome *K* is illustrated. *K* can be any integer value between 1 and 22.

### Extracting 3D genome-aware epigenetic features from BDM

After the BDM construction is completed, normal/stem references are constructed from the first principal component (PC1) of BDMs (Fig. S2B). Assuming that there are *n* samples belonging to the same tissue type (e.g., kidney), BDM of chromosome 1 (chr1) can be computed from each sample. The resulting *n* PC1 values are averaged to yield a normal reference for chr1 in kidney. Stem reference is computed by replacing the normal samples with stem cells. Afterwards, the normal/stem distances are computed by measuring the distance between PC1 vectors (Fig. S2C). In detail, the distance from sample’s PC1 to the corresponding normal/stem reference is measured for each autosome. Consequently, total 44 values are measured, each half of them originating from the use of normal and stem references, respectively. The 22 values computed by using the normal references are averaged to a single scalar value, or normal distance. Stem distance is examined alike, by utilizing values calculated between sample’s PC1 and stem reference. Finally, stem closeness of each sample is evaluated using the stem/normal distances (Fig. S2D). Stem/normal references are plotted into a cartesian 2D space, and a straight line connecting the origin and a dot representing each sample is drawn. The angle (*θ*) between this line and the *x*-axis is computed, and cos *θ* is used as stem closeness.

The procedure of measuring stem closeness is based on the observation that significant positive correlation exists between normal and stem distances from all cohorts used in the experiment, and the hypothesis that if stem and normal distances always move in the same direction, the smaller increment of stem distance followed by a unit increase (e.g., +0.1) of normal distance would imply higher stem-likeness of a given sample compared to other samples. The usage of cos *θ* is based on this rationale, along with some background concepts: (1) in the cartesian 2D space whose *x* and *y* axes represent the normal and stem distances, respectively, all samples’ points are in the first quadrant since the distances are nonnegative, (2) the angle between *x* axis and any arbitrary point located in the first quadrant (measured in degree) is in the range of [0, 90°], and (3) in this range of angle, cos *θ* keeps decreasing as *θ* increases. Therefore, samples in the steeper line would have smaller cos *θ* values compared to those in the line with gentler slope. This idea is tested on various cancer types, and it is suggested that the results were consistent with our assumption. In detail, the normal samples had smaller cos *θ* values compared to the tumor samples, thereby located closer to the *y* axis. The most distinct example of this, the one with TCGA-KIRP, is illustrated in Fig. S2D.

### Figuring out the optimal stem closeness for each cancer type

In order to determine the optimal stem closeness for each type of cancer, all possible combinations of different parameters were examined. After careful analysis, the best working combination for each cohort was identified. The finalized combination of parameters for each cohort are described in Table S9. Each subsection explains the parameters used in the experiment, and the method for determining the optimal combination of parameters, respectively.

#### Parameters

- Distance metric: The metric to be used when computing the distance between two PC1 vectors. Possible options are euclidean distance and cosine similarity. In case of the latter, a pseudocount of 10*^−^*^1^^5^ was added to the similarity, and an inverse of the resulting value was used as a distance.
- Matrix type: Interestingly, from preliminary experiments, it was found that PC1s from the inverse exponential of BDM (IEBDM) have remarkably high correlation with BDM PC1s. Based on this observation, IEBDM PC1s were also utilized. When using IEBDM, BDM was replaced with IEBDM throughout the whole pipeline illustrated in Fig. S2.
- Averaging method: To measure the stem/normal distances, 22 distances between BDM PC1s from each individual and corresponding reference PC1s are averaged. In this step, either the simple or weighted averaging was used. For the latter case the ratio of each autosome length to the sum of all autosome lengths are used as weights.
- Min-max scaling: If this option is used, normal/stem distances were min-max scaled into the range of [0, 1]. After recording the max and min values of normal/normal distances per cohort, each distance value was scaled according to Eq. S1. In Eq S1, *x_i_* denotes the normal/stem distance of the *i*-th sample, *min*(**x**) is the min value of normal/stem distances of current cohort, and *max*(**x**) is the max value of normal/stem distances of current cohort. This procedure was conducted between the steps depicted in Fig. S2C and Fig. S2D.

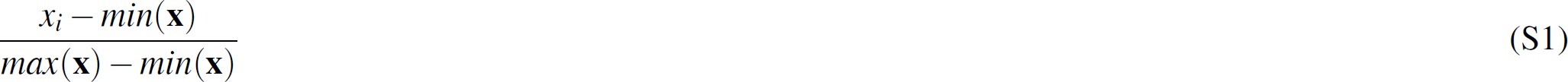
- Normalization: If this option is used, normal/stem distances were scaled into the range of [0, 1] according to Eq. S2. In Eq. S2, *x_i_* represents the normal/stem distance of the *i*-th sample, and *max*(**x**) is the max value of normal/stem distances of current cohort.

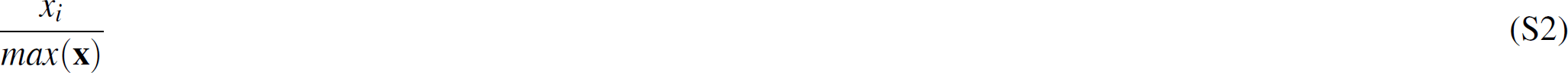
- Standardization: If this option is used, PC1 vectors were standardized according to Eq. S3 before computing distance. In Eq. S3, **y** represents each PC1 vector, *y _j_* denotes the *j*-th entry of **y**, *mean*(**y**) is the average value of all entries in **y**, and *std*(**y**) is the standard deviation of all entries in **y**.

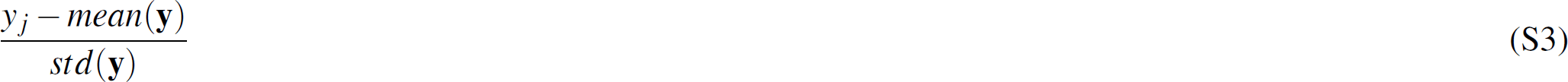
- Number of chromosomes:^9^ demonstrated that using smaller chromosomes lead to the declined performance of 3D genome reproduction and utilizing the entire chromosomes sometimes performed worse compared to the case of using only a part of genomic bins. Based on these observations, the number of autosomes used was set as a parameter. In detail, PC1 from chr1 up to chr*n* was used (*n* is an integer in the range of [1, 22]). In Fig. S2C, the case of of *n* = 22 is depicted.

### Selecting one optimal score per cohort

After conducting the log-rank tests using stem closeness scores from each combination of parameters, the stem closeness scores were first grouped based on the number of survival events in which each score acted as a significant predictor (*m*). Since the log-rank test was carried out for the four survival events in total (Overall survival; OS, Disease-specific survival; DSS, Disease-free interval; DFI, and the Progression-free interval; PFI), *m* is an integer in the range of [0, 4]. Subsequently, the stem closeness included in the group of *m* = 4 were probed first. If no score existed in the current group-of-interest, *m* was decreased by 1. From the current group-of-interest, the sum of p-values (*sum*_*p*) was computed from the results of log-rank test where the stem closeness was a significant predictor. After computing *sum*_*p* of all stem closeness scores, the scores by which the high score group was predicted to have better prognosis compared to the low score group, which is the opposite result desired by the score, were excluded. Finally, the remaining scores for each cohort were ranked by *sum*_*p* in an increasing order, and the one with the smallest *sum*_*p* was selected as the final score. All results of the log-rank tests reported in this manuscript are based on the stem closeness scores selected according to these procedures. The same scores were also used for the Cox regression, following the log-rank test.

### Hi-C data processing

To process Hi-C data, the 4DN Hi-C processing pipeline was used^36^. Raw Hi-C sequencing data as fastq files for cancer^37^, normal^20^, and stem cell lines^38, 39^ were downloaded from Sequence Read Archive (SRA) using sra-tools (v2.10.1) and parallel-fastq-dump (hepatocellular carcinoma cell line: SRS2627396; colon cancer cell line: SRS3816279; breast cancer cell line: SRS3505364; esophageal adenocarcinoma cell line: SRS3505365; lung: SRS1704412 and SRS1704413; pancreas: SRS1704415, SRS1704416, SRS1704417, and SRS1704418; hESCs: SRS1688434 and SRS3533281). The sequencing reads are mapped on hg19 reference genome using bwa (v0.7.17)^40^. Afterwards, aligned files (bam) were converted and handled as files that represent Hi-C pair information by using pairtools (v.0.3.0)^41^ and Hi-C interaction frequency matrices were obtained using cooler^42^. Finally, A/B compartment analyses were performed using FAN-C^43^.

### Functional annotation of DMR genes

This section explains the detailed results of functional annotation on the DMR genes, the genes located in DMR defined by predicted risks. The detailed inspections on the significantly enriched Gene Ontology (GO) terms, excluding one on the RA signaling pathway which is elaborately explained in the main manuscript, are provided.

One noticeable result is that genes associated with the gas transport were found in DMRs. We conjectured this result to be related to hypoxia, a deficiency of oxygen, which frequently occurs in cancer^44, 45^. Interestingly, all DMR genes we found to be related to the gas transport were the genes encoding subunits of hemoglobin, which transports oxygen. One of these genes, *HBB*, is reported to be abnormally expressed in many cancer types^46–48^. This result suggests the possibility that genes frequently altered in cancer, although not directly related to the developmental process, can also be regulated by open sea CpG probes.

Furthermore, the terms related to cell adhesion, such as homophilic cell adhesion via plasma membrane adhesion molecules and cell-cell adhesion via plasma-membrane adhesion molecules, were enriched. Cell adhesion is a crucial factor for cancer progression. In cancers, cell adhesion molecules are abnormally altered, incurring the tumor cells to interconnect better with other cells. In detail, the increased interaction between cancer cells and endothelium leads to faster metastasis, causing worse prognosis^49, 50^.

Among the other enriched GO terms were the mitotic cell cycle. We assume this is related to the biological characteristics of cancer cells, considering that they arise from the defective cell cycles. In detail, cancer is caused by the malfunctioning mitotic cell cycle, in which the aberrations occur in the cell cycle itself (cell cycle checkpoints) or genes regulating the cell cycle (e.g., p53 and BRCA1 gene). As a result, the cell goes through uncontrolled growth^51–54^, progressing to cancer.

Finally, a GO term related to the gamma-aminobutyric acid (GABA) signaling pathway was among the significantly enriched terms. GABA is related to the development of many types of cells. Also, GABA is supposed to act as an important modulator across different cancer types. In addition, the increased level of GABA substantially enhances the cancer cells’ capacity to invade, implying the contributive role of GABA in metastasis. GABA receptors, along with GABA, are also known to regulate cell proliferation. Moreover, the gene expression of GABA receptors was related to cancer prognosis and tumorigenesis^55–58^.

All these results suggest that many validated cancer-related pathways were enriched in DMR genes. Considering that the open sea DNA methylation levels of DMR genes differ significantly between the risk-high and risk-low group, it can be posited that the altered DNA methylation levels of open sea CpG probes are involved in several cancer hallmarks. The overall results from functional analyses are provided in Table S8.

---

**Figure S1.**
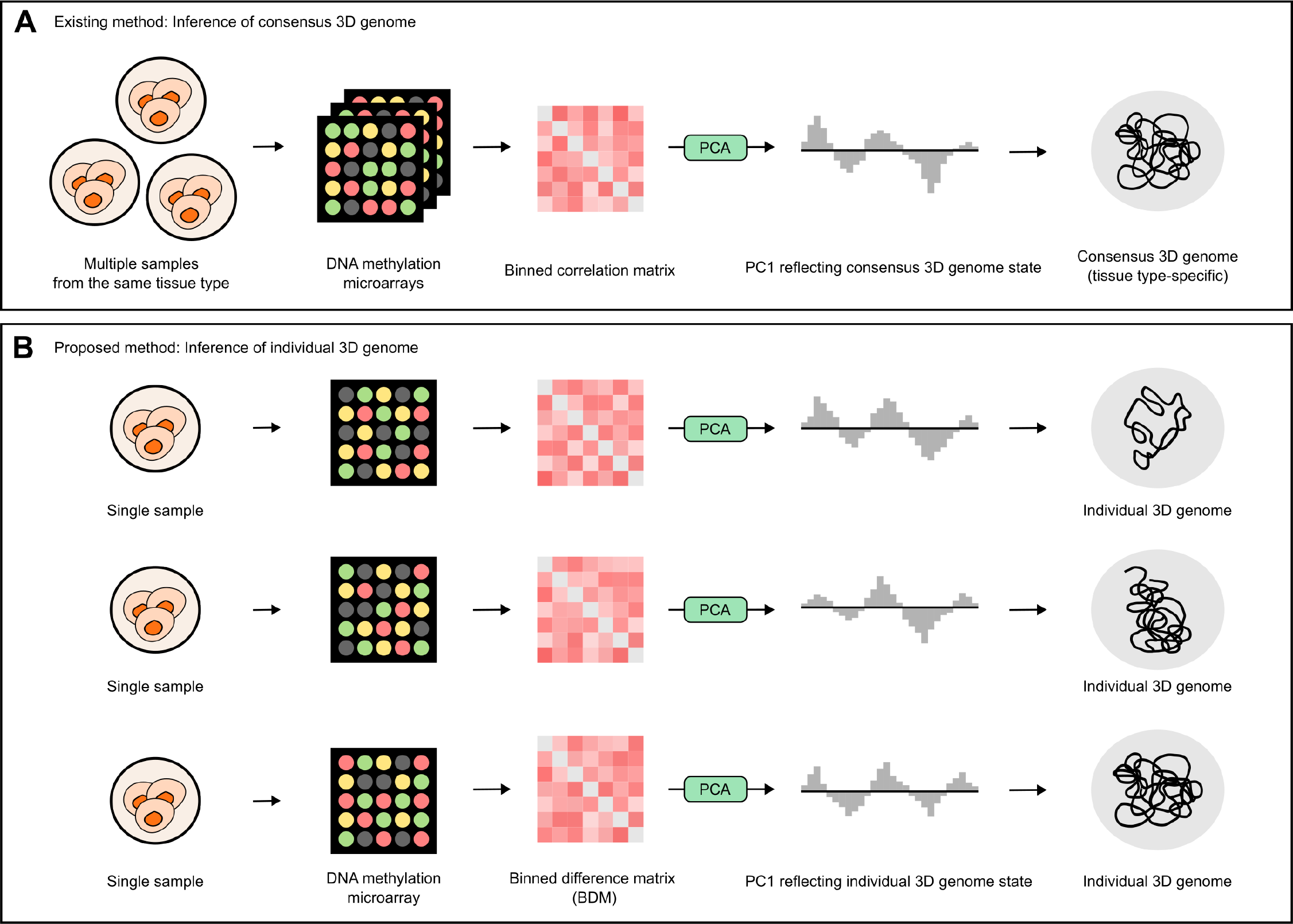
Comparison of existing and the proposed methods for 3D genome structure inference from the DNA methylation data. (A) Existing methods^9^. DNA methylation data of multiple samples from the same tissue type is used to infer a single consensus 3D genome structure. (B) Our proposed method. Single-sample based 3D genome structure inference from the DNA methylation data.

**Figure S2.**
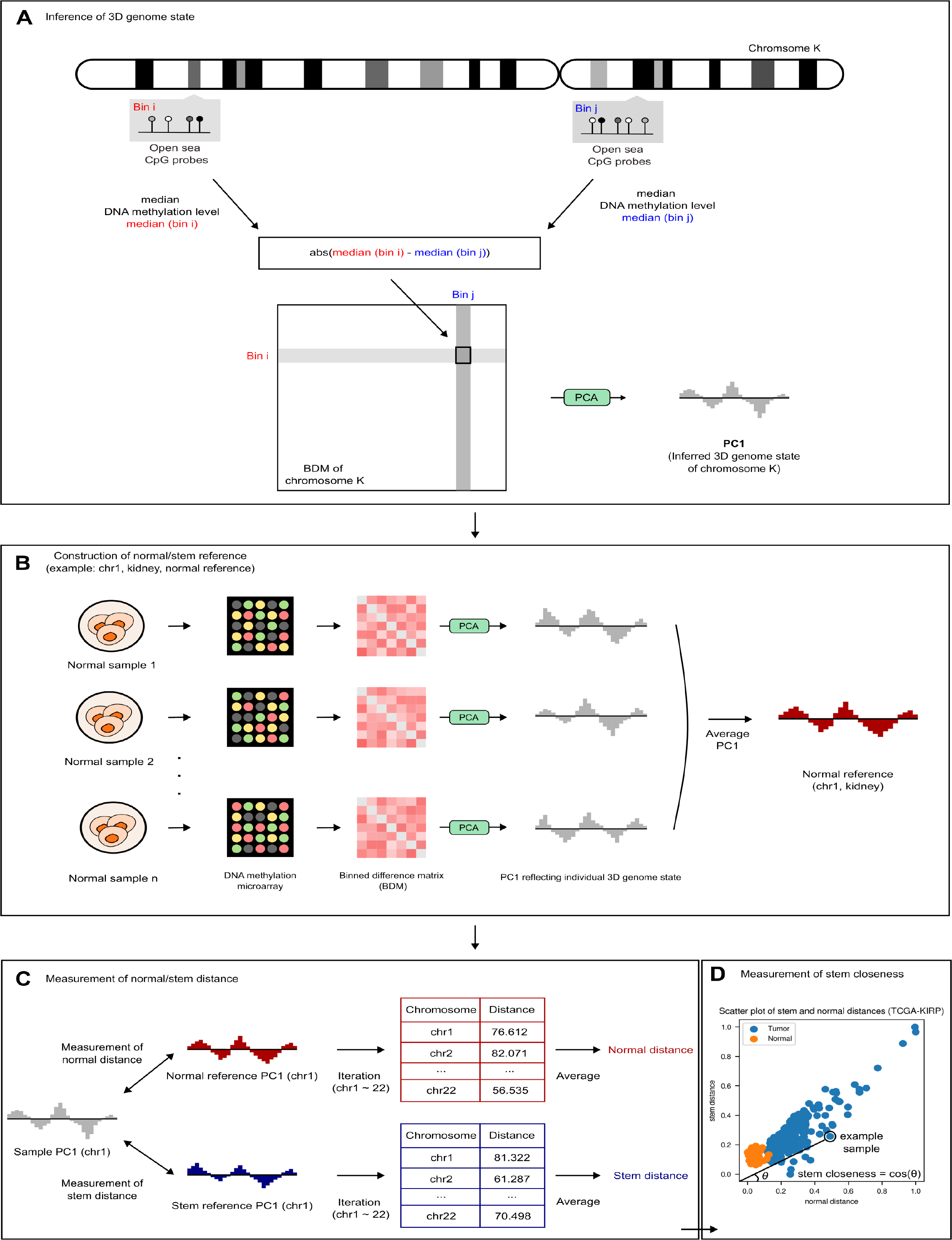
Quantitative explanation of the whole pipeline for extracting 3D genome-aware epigenetic features from DNA methylation data. (A) Inference of 3D genome structure from each autosome of a single sample. (B) Construction of normal/stem references. (C) Measurement of normal/stem distances. (D) Measurement of stem closeness.

**Figure S3.**
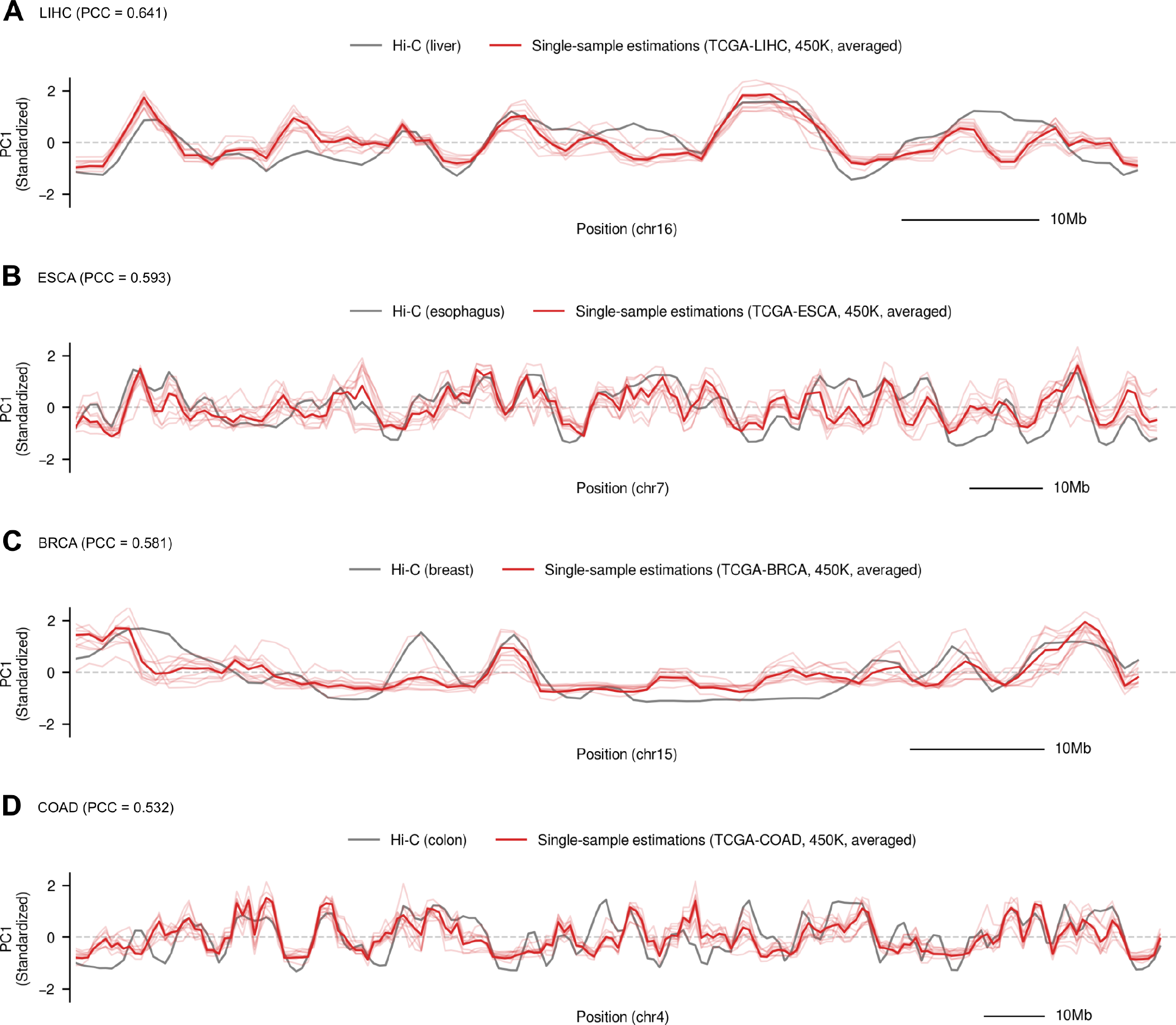
Comparison between the averaged BDM PC1 and Hi-C PC1s. The former and latter are extracted from TCGA tumor samples and the Hi-C data of cancer cell lines, respectively. Hi-C PC1s are plotted in gray. The averaged BDM PC1 and individual BDM PC1s used in averaging are plotted in dark red and light red, respectively. At the top left of each subplot, the TCGA cohort and PCC between averaged BDM PC1 and Hi-C PC1 is written. (A) LIHC tumor samples, liver hepatocellular carcinoma, chr16. (B) ESCA tumor samples, esophageal adenocarcinoma, chr7. (C) BRCA tumor samples, breast cancer, chr15. (D) COAD tumor samples, colon adenocarcinoma, chr4.

**Figure S4.**
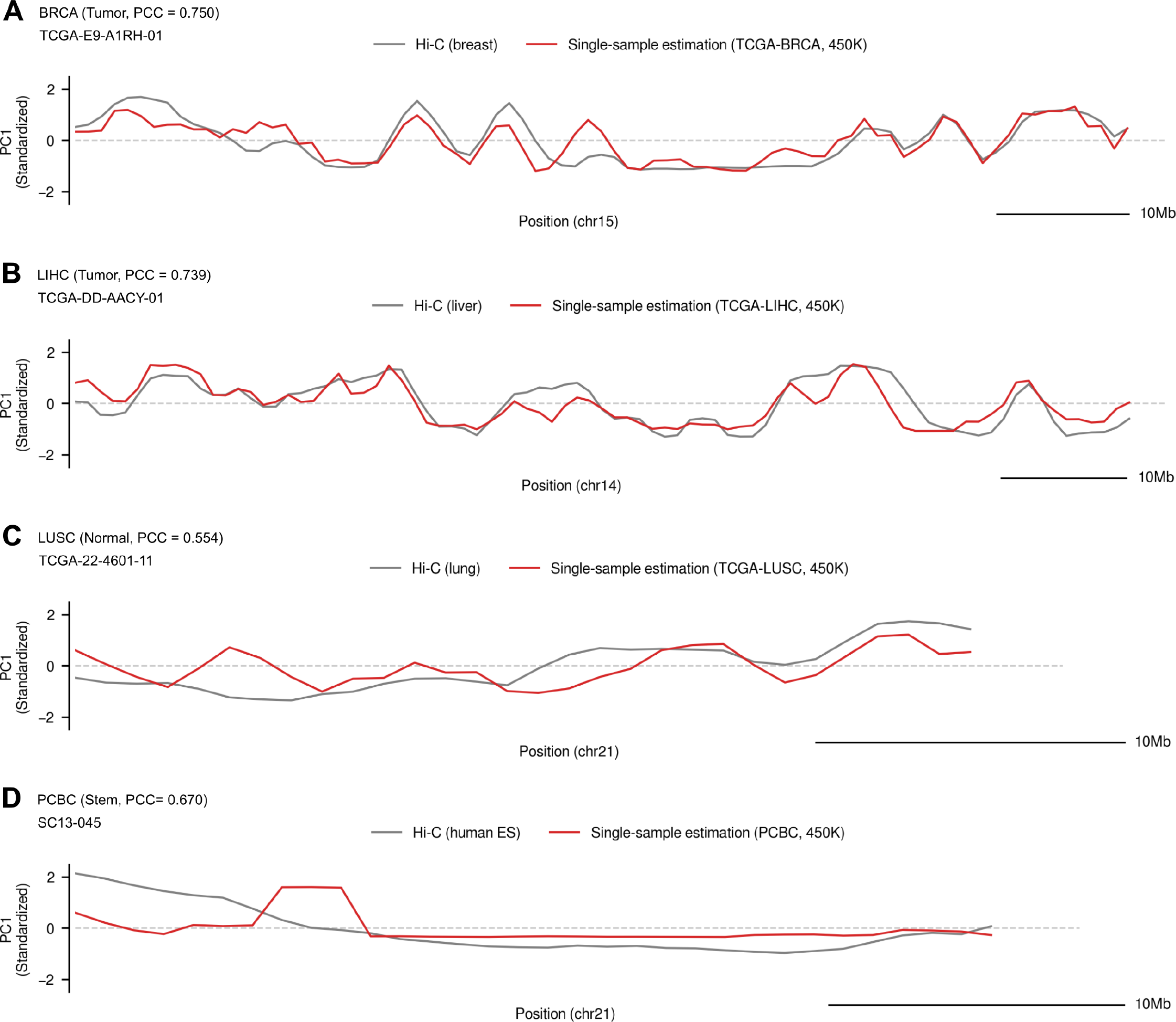
Comparison between the individual BDM PC1s and Hi-C PC1s. The formers are extracted from either the TCGA samples (tumor or normal samples) or stem cell samples (derived from the PCBC dataset^18^). The latters are extracted from Hi-C data of different cell lines, matched with the tissue type of BDM PC1s. Hi-C PC1s and the individual BDM PC1s are plotted in gray and dark red, respectively. At the top left of each subplot, the TCGA cohort (or ‘PCBC’ in case of stem cell samples) along with the category of tissue type (tumor, normal, or stem) and the PCC between BDM PC1 and Hi-C PC1. Name of TCGA or PCBC sample is written at the top left of each subplot, below the name of the cohort. (A) BRCA tumor samples, breast cancer cells, chr15. (B) LIHC tumor samples, liver hepatocellular carcinoma cells, chr14. (C) LUSC normal samples, normal lung tissue, chr21. (D) PCBC stem cells, H9 human embryonic stem cells, chr21

**Figure S5.**
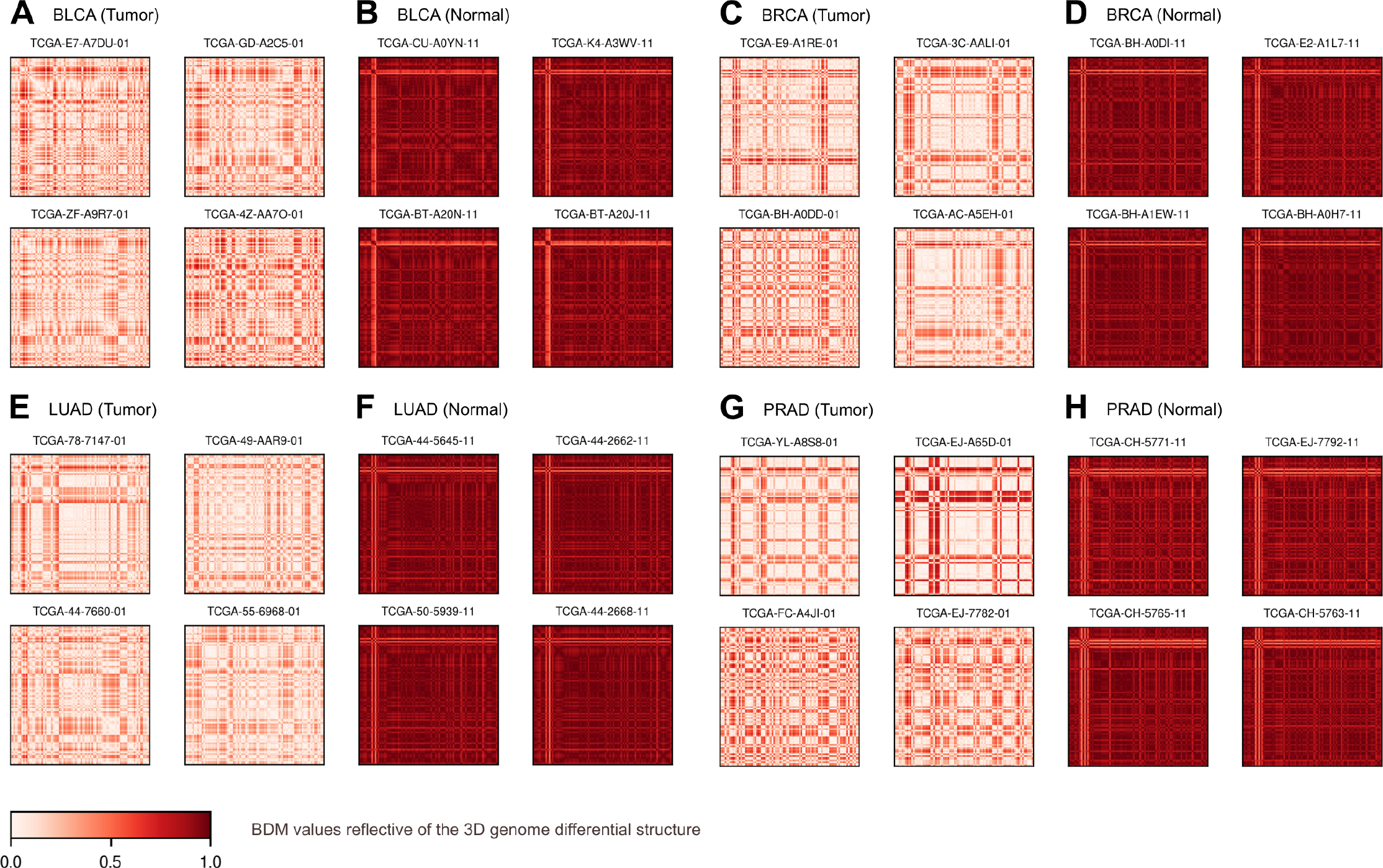
Heatmap visualization of the binned difference matrices (BDMs) from different TCGA cohorts. Entries of the BDM are in the range of [0, 1], as denoted by the colorbar located in the bottom left. At the top left of each subplot, the name of the TCGA cohort is written with the sample type (tumor or normal). For each subplot, BDMs of four samples are plotted. (A) BLCA, tumor samples. (B) BLCA, normal samples. (C) BRCA, tumor samples. (D) BRCA, normal samples. (E) LUAD, tumor samples. (F) LUAD, normal samples. (G) PRAD, tumor samples. (H) PRAD, normal samples.

**Figure S6.**
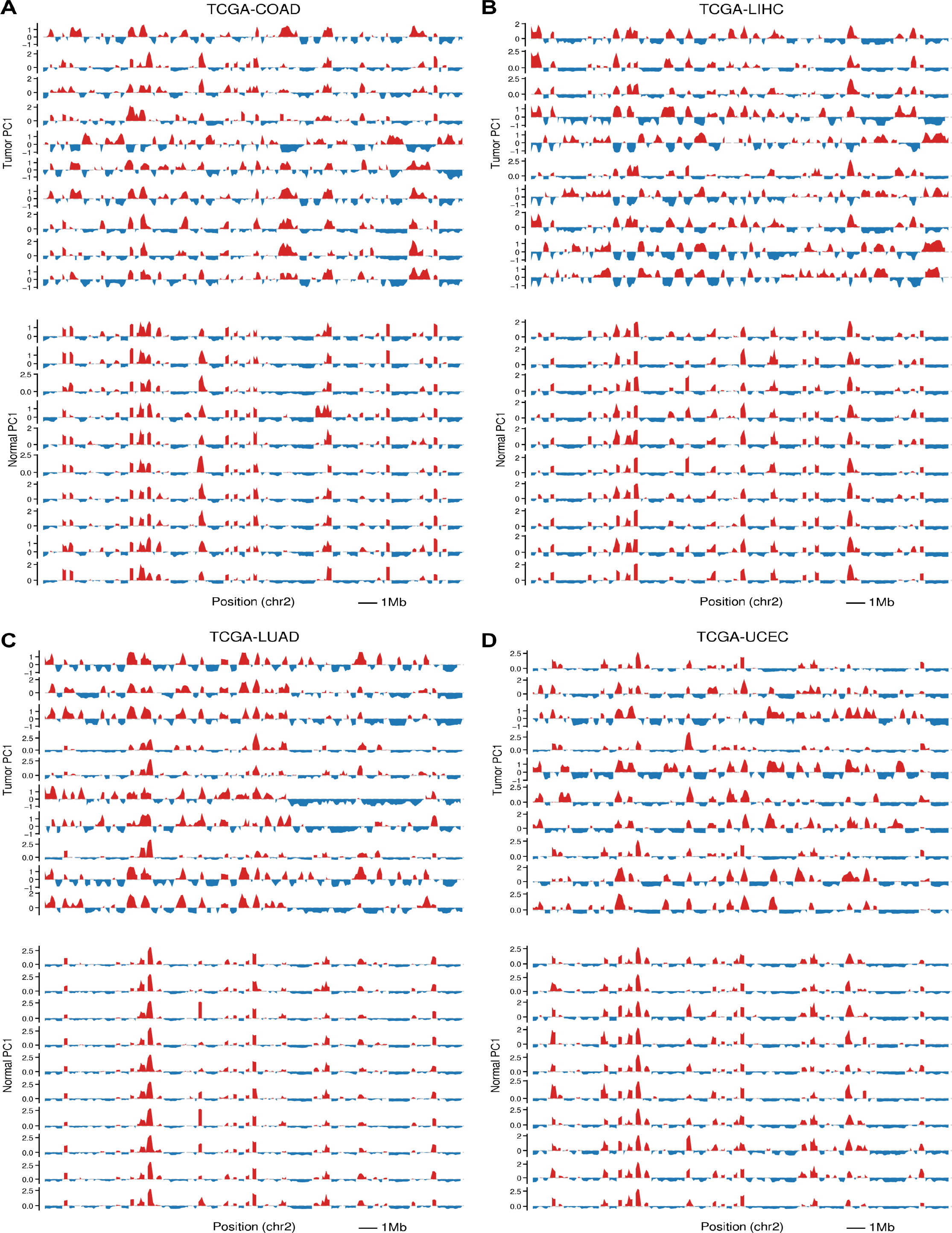
BDM PC1s of tumor and normal samples from different TCGA cohorts. The four cohorts are illustrated, in which the difference between BDM PC1s from the tumor and normal groups was the most distinct. (A) COAD, (B) LIHC, (C) LUAD, (D) UCEC.

**Figure S7.**
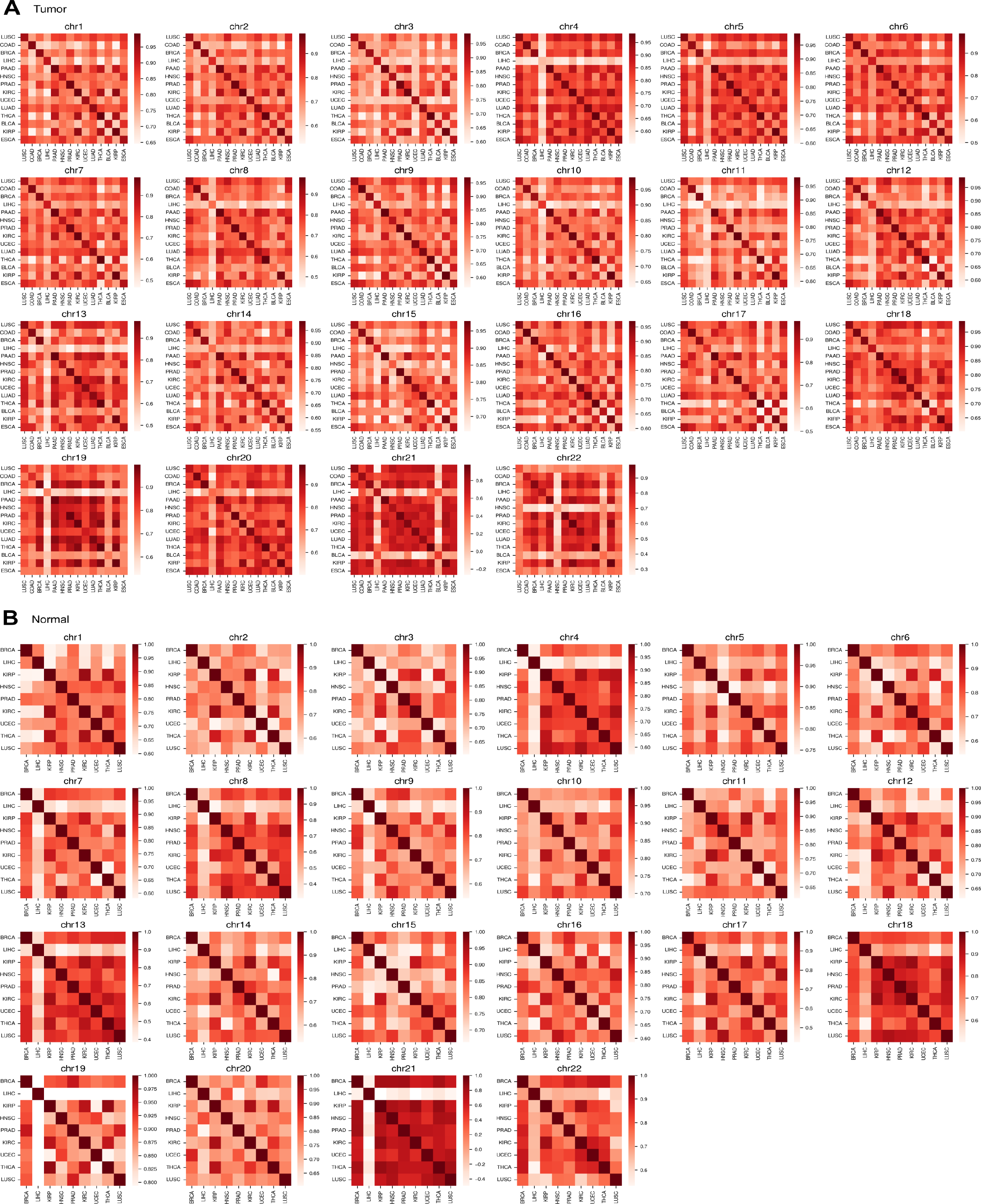
The Pearson correlation coefficient (PCC) between averaged BDM PC1s. The PCC values are visualized by heatmaps, separately for the tumor and normal groups of each TCGA cohort. (A) Heatmaps of PCC values from averaged BDM PC1s of tumor samples. (B) Heatmaps of PCC values from averaged BDM PC1s of normal samples. In both (A) and (B), the diagonal entries (the average PCC values for homogeneous pairs) have higher values than other entries.

**Figure S8.**
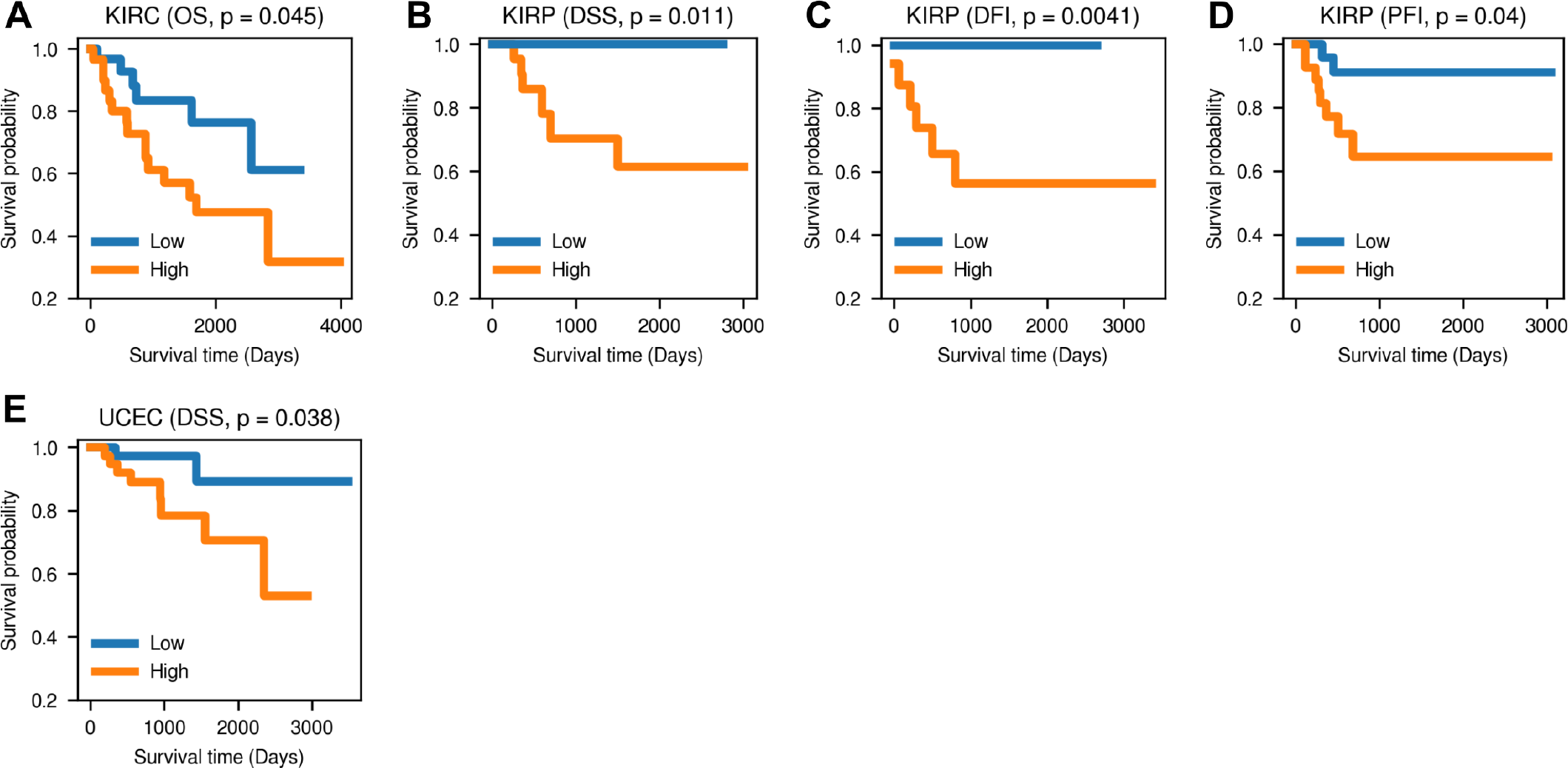
Result of the log-rank test, using risks predicted by feedforward neural networks as a predictor. OS, DSS, DFI, and PFI stands for the overall survival, disease-specific survival, disease-free interval, and the progression-free interval, respectively. The names of the cohort and survival event are written at the top center, followed by the log-rank test p-value. (A) KIRC (OS), (B) KIRP (DSS), (C) KIRP (DFI), (D) KIRP(PFI), and (E) UCEC (DSS).

**Figure S9.**
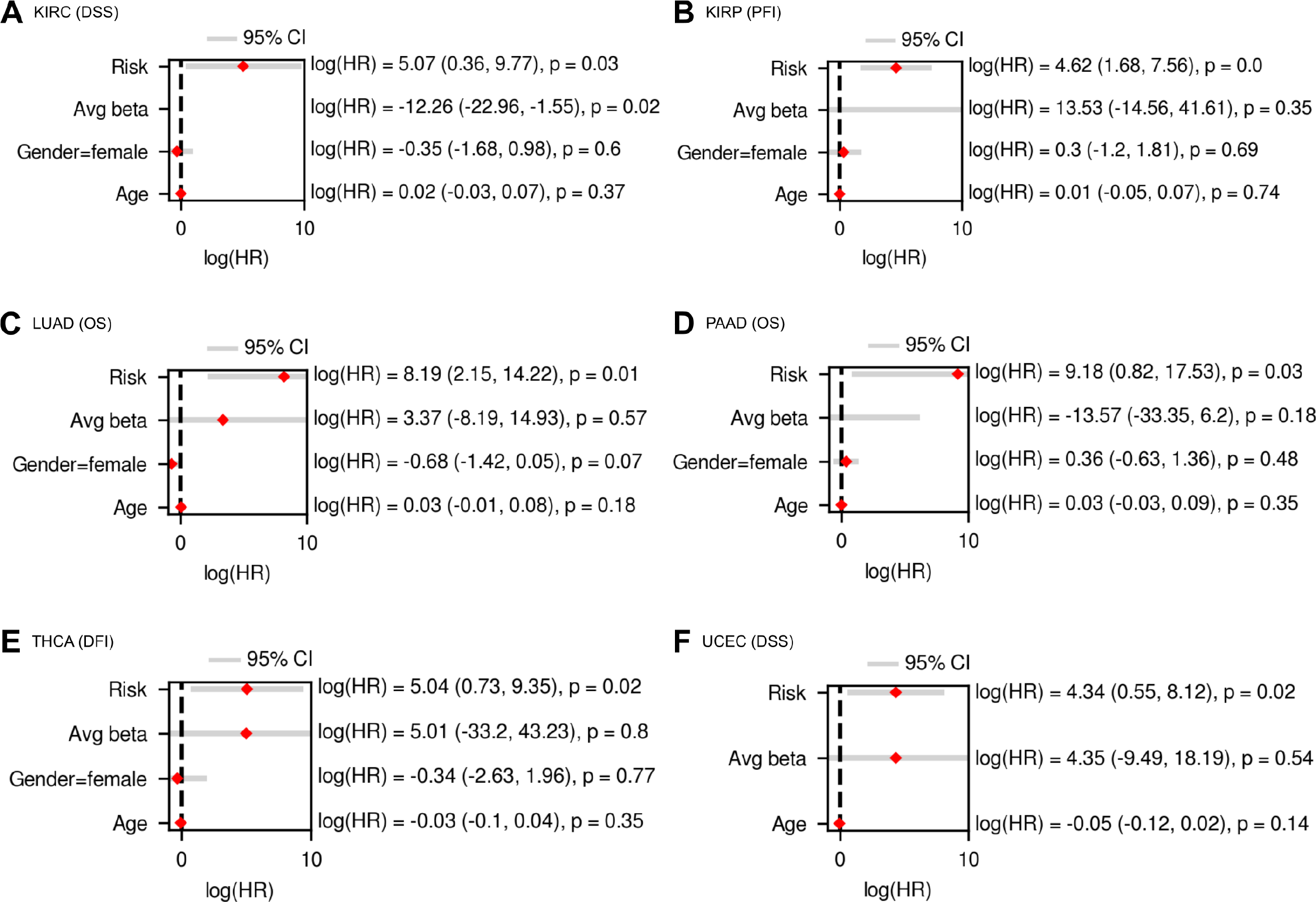
Cox regression results from the following input covariates: age, gender, predicted risk, and the average open sea DNA methylation level (the 3D genome-unaware epigenetic feature). Six cases in which predicted risk was a significant covariate are plotted. OS, DSS, DFI, and PFI stands for the overall survival, disease-specific survival, disease-free interval, and the progression-free interval, respectively. HR stands for hazard ratio. The black vertical line demarcates HR = 0. At the right side of each subplot, log(HR) of each input covariate is written, followed by the 95% confidence interval in the parenthesis and the p-value. (A) KIRC (DSS), (B) KIRP (PFI), (C) LUAD (OS), (D) PAAD (OS), (E) THCA (DFI), (F) UCEC (DSS)

**Figure S10.**
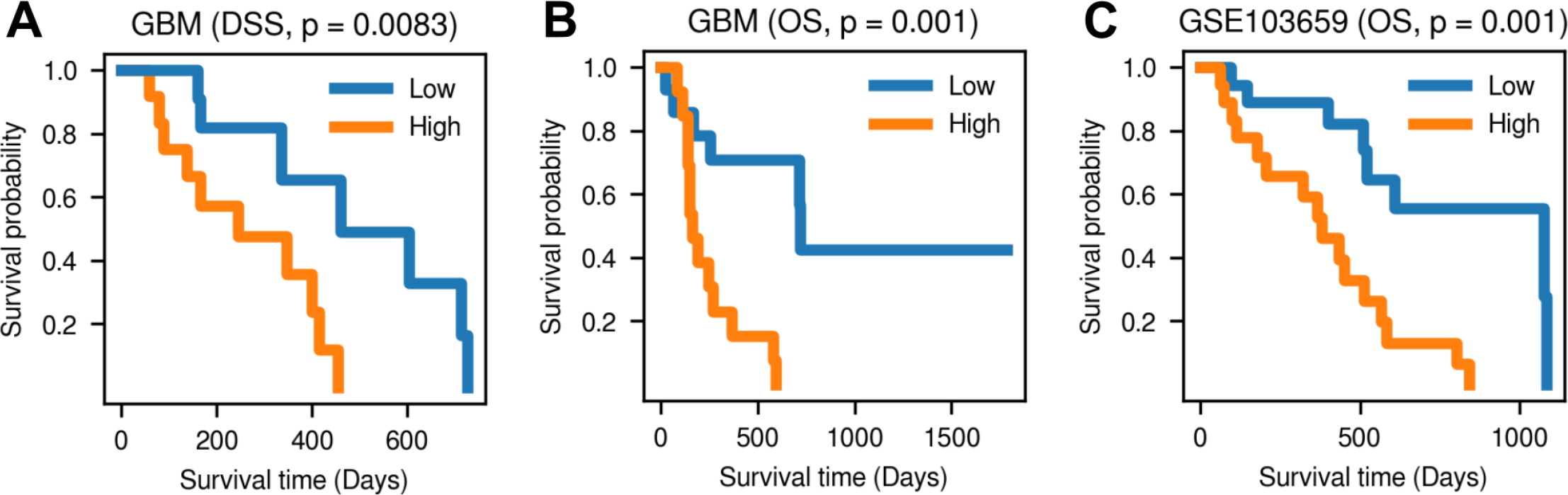
Significant log-rank test results from TCGA-GBM and GSE103659, using risk predicted from 3D genome-aware epigenetic features. OS, DSS, DFI, and PFI stands for the overall survival, disease-specific survival, disease-free interval, and the progression-free interval, respectively. The names of the cohort and survival event are written at the top center, followed by the log-rank test p-value. (A) GBM (DSS), (B) GBM (OS), (C) GSE103659 (OS)

**Figure S11.**
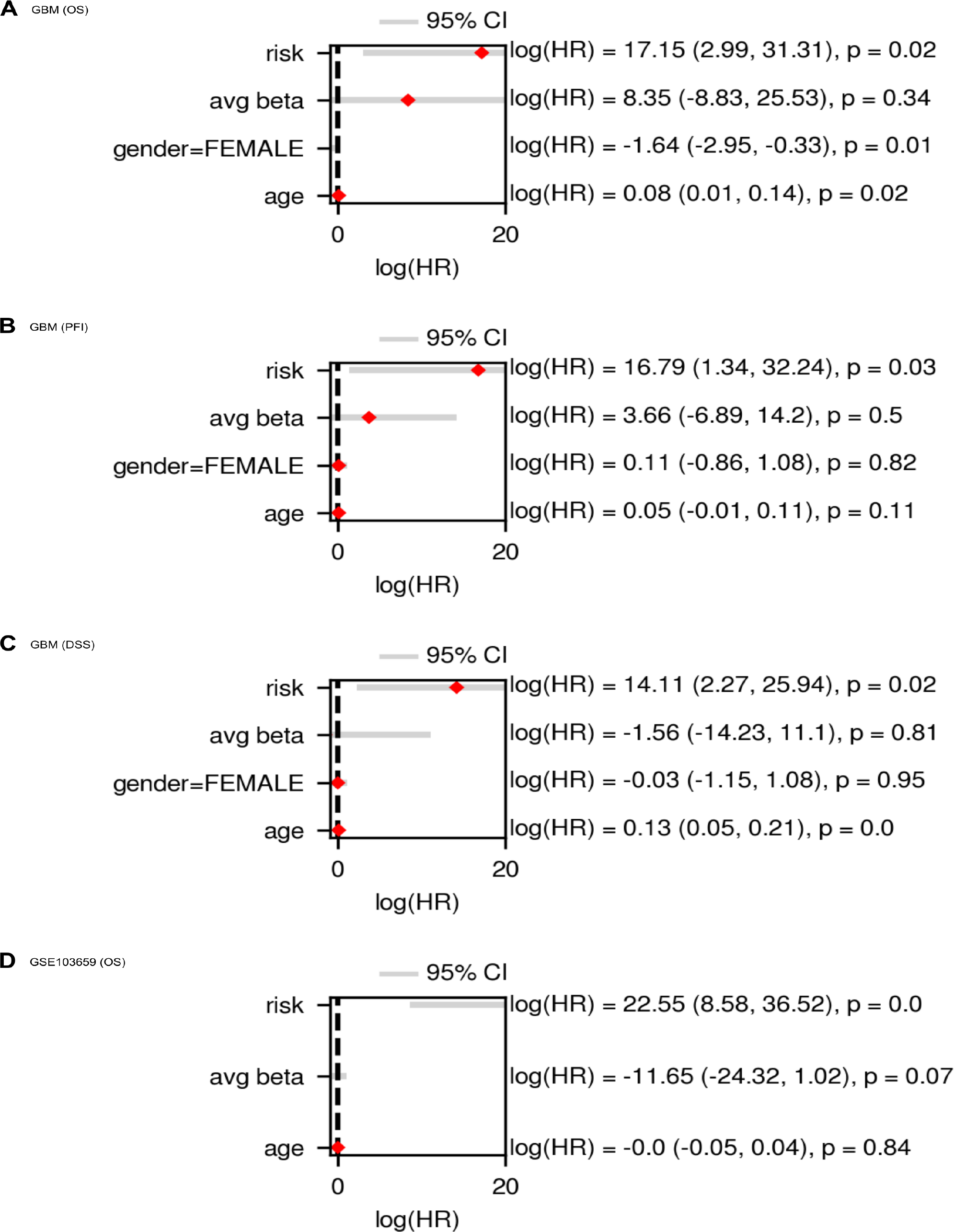
Cox regression results from TCGA-GBM and GSE103659, in which the predicted risk was a significant predictor. At the top left of each subplot, the names of the cohort and survival event are written. OS, DSS, DFI, and PFI stands for the overall survival, disease-specific survival, disease-free interval, and the progression-free interval, respectively. HR stands for hazard ratio. The black vertical line demarcates HR = 0. At the right side of each subplot, log(HR) of each input covariate is written, followed by the 95% confidence interval in the parenthesis and the p-value. (A) GBM (OS), (B) GBM (PFI), (C) GBM (DSS), (D) GSE103659 (OS)

**Figure S12.**
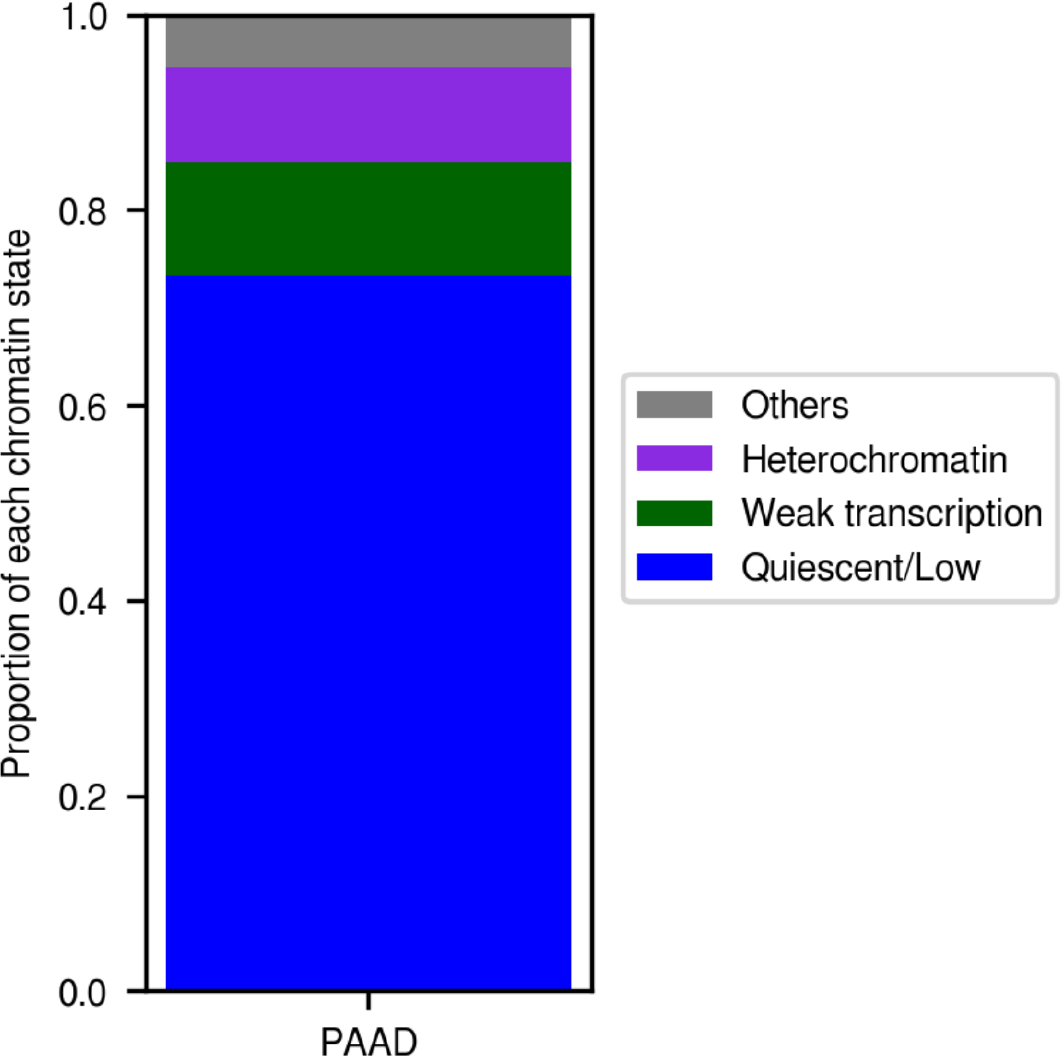
Proportion of the chromatin states in differentially methylated regions (DMR) defined by predicted risks. The *x* and *y* axes represent the name of TCGA cohort and proportions of different chromatin states, respectively. Among the three most dominant states, two (**Quiescent/Low** and **Heterochromatin**) are inactive states and the other one (**Weak transcription**) is a slightly activated state. All the other chromatin states excluding these three dominant states are labelled as ‘Others’. The results imply that genes located in regions with dominant chromatin states, which are not actively expressed in normal situation, could be anomalously influenced by the altered DNA methylation level in risk-high group.

**Table S1.** Description and composition of each TCGA cohort

| TCGA cohort | Description | n_tumor | n_normal | n_total |
| --- | --- | --- | --- | --- |
| BLCA | Bladder urothelial carcinoma | 413 | 21 | 434 |
| BRCA | Breast invasive carcinoma | 790 | 98 | 888 |
| CHOL | Cholangiocarcinoma | 36 | 9 | 45 |
| COAD | Colon adenocarcinoma | 299 | 38 | 337 |
| ESCA | Esophageal carcinoma | 186 | 16 | 202 |
| HNSC | Head and Neck squamous cell carcinoma | 530 | 50 | 580 |
| KIRC | Kidney renal clear cell carcinoma | 320 | 160 | 480 |
| KIRP | Kidney renal papillary cell carcinoma | 276 | 45 | 321 |
| LIHC | Liver hepatocellular carcinoma | 379 | 50 | 429 |
| LUAD | Lung adenocarcinoma | 460 | 32 | 492 |
| LUSC | Lung squamous cell carcinoma | 372 | 43 | 415 |
| PAAD | Pancreatic adenocarcinoma | 185 | 10 | 195 |
| PRAD | Prostate adenocarcinoma | 499 | 50 | 549 |
| THCA | Thyroid carcinoma | 515 | 56 | 571 |
| UCEC | Uterine Corpus Endometrial Carcinoma | 432 | 46 | 478 |

**Table S2.**
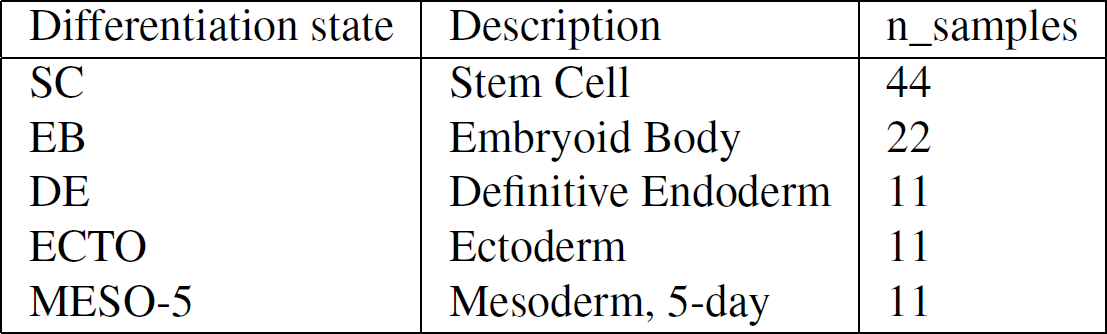
Composition of stem cell samples

| Differentiation state | Description | n_samples |
| --- | --- | --- |
| SC | Stem Cell | 44 |
| EB | Embryoid Body | 22 |
| DE | Definitive Endoderm | 11 |
| ECTO | Ectoderm | 11 |
| MESO-5 | Mesoderm, 5-day | 11 |

**Table S3.** TCGA cohorts matched to the Hi-C data of cancer cell lines^17, 19^.

| TCGA cohort | Hi-C cancer cell line | Hi-C cancer cell line description | GEO IDs of Hi-C data |
| --- | --- | --- | --- |
| BRCA | HCC1954 (Breast cancer cell line) | Breast_Cancer | GSM3258551 |
| COAD | SW480_ (Colon cancer cell line) | Colon_Adenocarcinoma | GSM3399745 |
| ESCA | OE33 (Esophageal adenocarcinoma cell line) | Esophageal_Adenocarcinoma | GSM3258552 |
| KIRC | G-401 (kidney cancer cell line) | Kidney_Renal_Clear_Cell_Carcinoma | GSM2825105, GSM2825106 |
| LIHC | HepG2 (hepatocellular carcinoma cell line) | Liver_Hepatocellular_carcinoma | GSM2825569, GSM2825570 |
| LUSC | NCI-H460 (lung cancer cell line) | Lung_Squamous_Cell_Carcinoma | GSM2827554, GSM2827555 |
| PAAD | Panc1 (pancreatic carcinoma cell line) | Pancreatic_Cancer | GSM2827313, GSM2827314 |
| PRAD | 22Rv1 (prostate cancer cell line) | Prostate_Adenocarcinoma | GSM3358191, GSM3358192 |

**Table S4.** TCGA cohorts matched to the Hi-C data of normal cell lines^17, 19^.

| TCGA cohort | Hi-C normal cell line | GEO IDs of Hi-C data |
| --- | --- | --- |
| PAAD | Pancreas | GSM2322547, GSM2322548, GSM2322549, GSM2322550 |
| LUSC | Lung | GSM2322544, GSM2322545 |
| LUAD | Lung | GSM2322544, GSM2322545 |

**Table S5.** PCBC stem cells matched to the Hi-C data of stem cell lines^18, 19^.

| Cohort | Cohort description | Hi-C stem cell line | GEO ID of Hi-C data |
| --- | --- | --- | --- |
| PCBC | Human embryonic stem cells | H9 Human Embryonic Stem Cells | GSM2309023 |
| PCBC | Human embryonic stem cells | Embryonic stem cell | GSM3263085 |

**Table S6.** Risk prediction results from the feedforward neural network. **[Top]** The results from the main scenario, using the concatenation of 3D genome-aware epigenetic features (BDM PC1, normal/stem distances, and stem closeness) and survival-related features (age and gender) as input to the neural network. [Middle] The results from baseline scenario 1 (BS1), where the risks were predicted from the age and gender without any epigenetic features. [Bottom] The results from baseline case 2 (BS2), where the risks were predicted from age, gender, and the 3D genome-unaware epigenetic feature (the average open sea DNA methylation level). The best performances are highlighted in bold.

| Cohort | Test set c-index | Log-rank p-value | Event |
| --- | --- | --- | --- |
| CHOL | 0.750 | 0.029 | DSS |
| KIRC | 0.742 | 0.049 | DSS |
| KIRC | 0.705 | 0.045 | OS |
| KIRP | 0.841 | **0.001** | OS |
| KIRP | **0.959** | 0.024 | DSS |
| KIRP | 0.902 | 0.004 | DFI |
| KIRP | 0.839 | 0.040 | PFI |
| PAAD | 0.688 | 0.016 | PFI |
| PRAD | 0.739 | 0.026 | DFI |
| THCA | 0.742 | 0.028 | OS |
| UCEC | 0.654 | 0.038 | DSS |
| UCEC | 0.669 | 0.042 | DFI |
| KIRC (**BS 1**) | 0.708 | **0.001** | OS |
| KIRP (**BS 1**) | 0.741 | 0.028 | DSS |
| PAAD (**BS 1**) | 0.810 | 0.026 | DFI |
| PRAD (**BS 1**) | 0.676 | 0.011 | DFI |
| THCA (**BS 1**) | **0.987** | 0.019 | OS |
| KIRC (**BS 2**) | 0.697 | **0.004** | OS |
| THCA (**BS 2**) | **0.929** | 0.024 | OS |
| THCA (**BS 2**) | 0.884 | 0.019 | OS |

**Table S7.** Risk prediction performance from TCGA-GBM and GSE103659. The bold texts represent the best performance (the biggest c-index and the smallest p-value).

| Cohort | Test set c-index | Log-rank p-value | Event |
| --- | --- | --- | --- |
| GBM | **0.743** | 0.008 | DSS |
| GBM | 0.733 | **0.001** | OS |
| GSE103659 | 0.721 | **0.001** | OS |

**Table S8.**
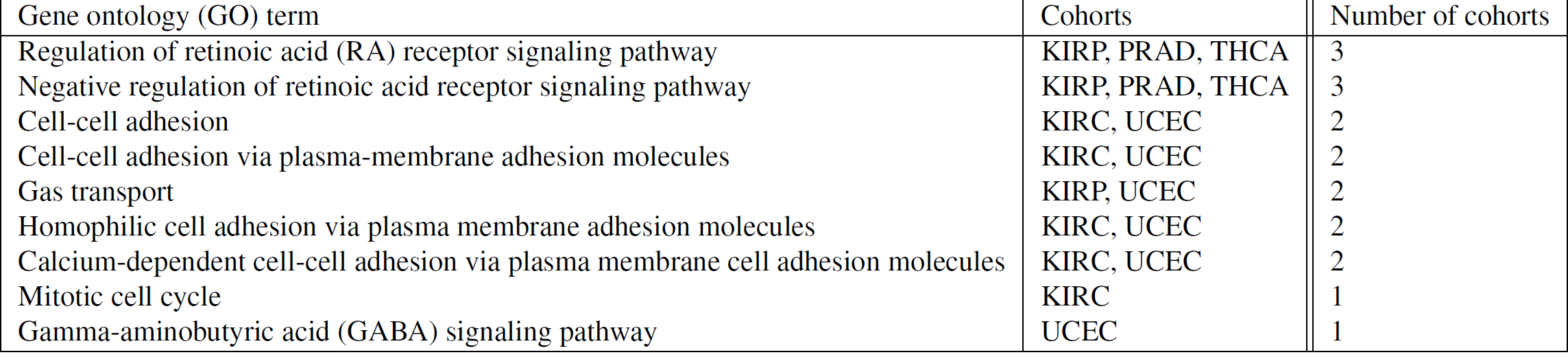
Significantly enriched terms found from DMR genes

| Gene ontology (GO) term | Cohorts | Number of cohorts |
| --- | --- | --- |
| Regulation of retinoic acid (RA) receptor signaling pathway | KIRP, PRAD, THCA | 3 |
| Negative regulation of retinoic acid receptor signaling pathway | KIRP, PRAD, THCA | 3 |
| Cell-cell adhesion | KIRC, UCEC | 2 |
| Cell-cell adhesion via plasma-membrane adhesion molecules | KIRC, UCEC | 2 |
| Gas transport | KIRP, UCEC | 2 |
| Homophilic cell adhesion via plasma membrane adhesion molecules | KIRC, UCEC | 2 |
| Calcium-dependent cell-cell adhesion via plasma membrane cell adhesion molecules | KIRC, UCEC | 2 |
| Mitotic cell cycle | KIRC | 1 |
| Gamma-aminobutyric acid (GABA) signaling pathway | UCEC | 1 |

**Table S9.** Finalized combinations of parameters for computing stem closeness

| Cohort | Distance metric | Matrix type | Averaging method | Minmax scaling | Normalization | Standardization | Number of chromosomes |
| --- | --- | --- | --- | --- | --- | --- | --- |
| BLCA | Cosine similarity | BDM | Simple average | Used | Not used | Used | 8 |
| BRCA | Cosine similarity | BDM | Simple average | Used | Not used | Not used | 7 |
| CHOL | Euclidean distance | BDM | Simple average | Not used | Used | Used | 19 |
| COAD | Euclidean distance | BDM | Simple average | Used | Not used | Used | 20 |
| KIRC | Euclidean distance | BDM | Weighted average | Not used | Used | Not used | 3 |
| KIRP | Euclidean distance | BDM | Weighted average | Used | Not used | Not used | 1 |
| LIHC | Cosine similarity | IEBDM | Simple average | Used | Not used | Used | 9 |
| LUAD | Euclidean distance | BDM | Weighted average | Not used | Used | Used | 1 |
| LUSC | Euclidean distance | BDM | Simple average | Used | Not used | Used | 14 |
| PAAD | Cosine similarity | BDM | Simple average | Used | Not used | Not used | 8 |
| PRAD | Euclidean distance | BDM | Simple average | Used | Not used | Used | 2 |
| THCA | Euclidean distance | BDM | Weighted average | Used | Not used | Used | 2 |
| UCEC | Euclidean distance | BDM | Simple average | Used | Not used | Used | 18 |

